# Myosin XVA isoforms participate in the mechanotransduction-dependent remodeling of the actin cytoskeleton in auditory stereocilia

**DOI:** 10.1101/2024.09.04.611210

**Authors:** Ana I. López-Porras, Ava M. Kruse, Mark T. McClendon, A Catalina Vélez-Ortega

## Abstract

Auditory hair cells form precise and sensitive staircase-like actin protrusions known as stereocilia. These specialized microvilli detect deflections induced by sound through the activation of mechano-electrical transduction (MET) channels located at their tips. At rest, a small MET channel current results in a constant calcium influx, which regulates the morphology of the actin cytoskeleton in the shorter ‘transducing’ stereocilia. However, the molecular mechanisms involved in this novel type of activity-driven plasticity in the stereocilium cytoskeleton are currently unknown. Here, we tested the contribution of myosin XVA (MYO15A) isoforms. We used electron microscopy to evaluate morphological changes in the cytoskeleton of auditory hair cell stereocilia after the pharmacological blockage of resting MET currents in cochlear explants from mice that lacked one or all isoforms of MYO15A. Hair cells lacking functional MYO15A isoforms did not exhibit MET-dependent remodeling in their stereocilia cytoskeleton. In contrast, hair cells that only lack the long isoform of MYO15A exhibited increased MET-dependent stereocilia remodeling, including remodeling in stereocilia from the tallest ‘non-transducing’ row of the bundle. We conclude that MYO15A isoforms not only enable but also fine-tune the MET-dependent remodeling of the actin cytoskeleton in transducing stereocilia and contribute to the stability of the tallest row.

## 1 Introduction

Inner ear hair cells have modified microvilli, known as stereocilia, on their apical surfaces that form highly organized bundles with rows of increasing height in a staircase manner. The cytoskeleton within each stereocilium contains hundreds of parallel and polarized actin filaments that are highly crosslinked, resulting in a rather rigid paracrystalline arrangement (Tilney et al., 1980, Zheng et al., 2000, Shin et al., 2010, Taylor et al., 2015). The tips of shorter stereocilia are linked to the shafts of their taller neighbors via extracellular connections called tip links, thus stereocilia bundle deflections in the direction of the tallest row increase the tension at the tips of shorter stereocilia (Pickles et al., 1984, Assad et al., 1991) where mechano-electrical transduction (MET) channels are located (Beurg et al., 2009). MET channels are non-selective cation channels (Corey and Hudspeth, 1979, Kros et al., 1992). To maintain the sensitivity of the hair bundle, tip links are kept under tension even at rest, which results in the constant entry of calcium into stereocilia (Hudspeth and Corey, 1977, Kros et al., 1992). We and others previously showed that this resting calcium entry impacts the morphology of the stereocilia cytoskeleton. In an earlier study, we used MET channel blockers, disruption of tips links, and manipulations of intracellular and extracellular calcium to show that a decrease in the resting calcium entry leads to the thinning of stereocilia tips and their eventual shortening (Velez-Ortega et al., 2017). Conversely, an increase in resting calcium entry leads to thickening of the stereocilia tips while removal of MET channel blockers or tip link regeneration induces stereocilia regrowth (Velez-Ortega et al., 2017). During the postnatal development of the hair bundle in the mouse, MET currents (and/or components of the MET machinery) are essential to establish correct stereocilia dimensions (height and thickness) in all rows of the bundle, and for the proper trafficking of certain stereocilia tip proteins that are differentially expressed between the tallest and the shorter stereocilia (collectively known as ‘row-identity’ proteins) (Krey et al., 2020, Krey et al., 2023). Altogether, these findings highlight a clear interplay between MET activity and stereocilia cytoskeleton remodeling. However, the exact molecular machinery involved in this process is still unknown. This study explores whether MYO15A, the master protein regulating stereocilia elongation during hair cell bundle development, is involved in the MET-dependent remodeling of the actin cytoskeleton within mammalian auditory stereocilia.

MYO15A is a non-conventional myosin that localizes to the tips of stereocilia (Belyantseva et al., 2003). It delivers proteins required for the proper elongation of the stereocilium actin core, such as the scaffolding protein WHRN (Belyantseva et al., 2005), the actin capping/bundling protein EPS8 (Manor et al., 2011), and the GPSM2-GNAI3 signaling complex (Tadenev et al., 2019). Functional defects in MYO15A, WHRN, EPS8, GPSM2 or GNAI3 all lead to profound hearing loss and auditory hair cells with abnormally short stereocilia (Probst et al., 1998, Mburu et al., 2003, Manor et al., 2011, Tadenev et al., 2019). Three isoforms of MYO15A have been identified in the auditory hair cells (Liang et al., 1999, Belyantseva et al., 2003, Rehman et al., 2016, Ranum et al., 2019), but only two of them have been characterized in detail in the mouse. Isoform 2 is the shortest, its mRNA levels are high at birth and quickly decline afterwards, and the protein is preferentially trafficked to the tips of stereocilia from the tallest row of the bundle. Isoform 1 is the longest isoform due to the addition of a large (∼130 kDa) N-terminal domain, its mRNA levels continually increase during the first postnatal weeks, and the protein is preferentially trafficked to the tips of stereocilia from the shorter rows of the bundle. Isoform 3 has a unique small N-terminal domain making it intermediate in size, but its localization within the hair cell bundle remains to be characterized.

In *shaker-2* mice (*Myo15^sh2/sh2^*), a point mutation in the motor domain of MYO15A renders all isoforms non-functional (*i.e*. unable to reach the stereocilia tips) (Belyantseva et al., 2003) and while stereocilia are abnormally short (Probst et al., 1998) they still exhibit MET currents (Stepanyan et al., 2006, Stepanyan and Frolenkov, 2009). In mice lacking the long isoform of MYO15A only (*Myo15^ΔN/ΔN^*), auditory hair cells develop normally, exhibit nearly normal MET currents, but their shorter (‘transducing’) stereocilia degenerate prematurely (Fang et al., 2015). Here, we evaluated the MET-dependent remodeling of the stereocilia cytoskeleton in auditory hair cells from *Myo15^sh2/sh2^* and *Myo15^ΔN/ΔN^* mice. Our results indicate that MYO15A is essential to deliver the machinery involved in MET-dependent stereocilia remodeling. In addition, the long isoform of MYO15A also delivers machinery that provides cytoskeleton stability.

## 2 Results

### 2.1 MYO15A is required for the MET-dependent remodeling on the stereocilia cytoskeleton

We previously showed that the blockage of the resting MET current with pharmacological MET channel blockers like tubocurarine (Farris et al., 2004) or benzamil (Rusch et al., 1994) induces the remodeling of the stereocilium cytoskeleton in auditory hair cells (Velez-Ortega et al., 2017). To test the role of MYO15A in this MET-dependent remodeling of stereocilia we used *Shaker-2* mice which have a point mutation in the motor domain of MYO15A that renders all MYO15A isoforms unable to climb the stereocilia tips. While the *shaker-2* hair cell bundles are abnormally short, *shaker-2* auditory hair cells still exhibit MET currents, including currents at resting bundle positions (Stepanyan et al., 2006, Stepanyan and Frolenkov, 2009). Therefore, we used MET channel blockage to evaluate the remodeling of neonate auditory stereocilia from heterozygous (*Myo15^sh2/+^*) and homozygous (*Myo15^sh2/sh2^*) *shaker-2* littermates.

As expected, 300 µM of tubocurarine significantly reduced the uptake of the FM1-43 dye through MET channels open at rest in inner and outer hair cells (IHC and OHC, respectively) from heterozygous and homozygous *shaker-2* mice (Fig. 1A). After 4 hours of MET channel blockage with 300 µM of tubocurarine, scanning electron microscopy (SEM) images showed significant thinning of second row (‘transducing’) stereocilia tips in postnatal day 5 (P5) heterozygous but not homozygous OHC (Fig. 1B,C). In addition to the stereocilia thinning, significant stereocilia shortening was observed in second row stereocilia from heterozygous but not homozygous mice (Fig. 1D). In our previous study (Velez-Ortega et al., 2017), we found that MET channel blockage does not affect the morphology of stereocilia from the tallest row in wild-type bundles, which do not express MET channels (Beurg et al., 2009). Similarly, here we found no signs of stereocilia shortening in OHC from either heterozygous or homozygous *shaker-2* mice (Fig. 1D).

**Figure 1.**
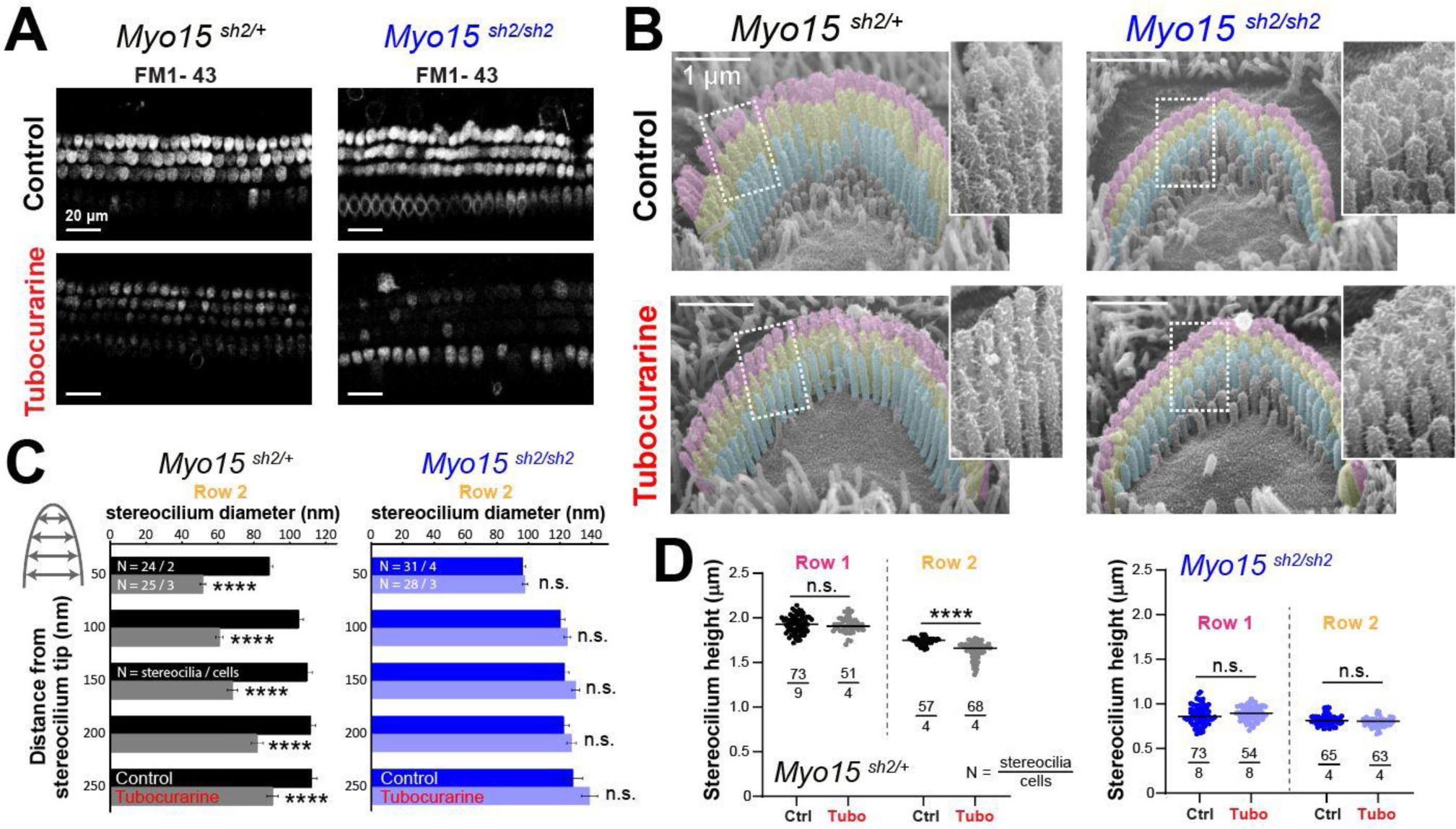
Outer hair cells lacking functional MYO15A do not exhibit mechanotransduction-dependent stereocilia remodeling. **(A)** Decreased FM1-43 dye uptake in hair cells after application of the MET channel blocker, tubocurarine (300 µM), in heterozygous (*left*) and homozygous *shaker2* (*right*) cochlear explants. **(B)** Representative false-colored scanning electron microscopy (SEM) images of outer hair cell (OHC) stereocilia bundles cultured for 4 hours at 37°C in control conditions (*top*) or with 300 µM tubocurarine (*bottom*), from heterozygous control (*left*) or homozygous *shaker2* (*right*) littermates. Insets show highlighted bundle regions at higher magnification. **(C)** Diameter of stereocilia from the second row (colored in yellow in panel B) at several positions from the stereocilium tip (as indicated in cartoon), from OHC of control heterozygous (black, gray) or homozygous *shaker2* (dark and light blue) littermates cultured in control conditions (darker bars) or with tubocurarine (lighter bars). Data are shown as mean ± SE. Statistical differences were analyzed using a linear mixed model. **(D)** Heights of stereocilia from the first and second rows (colored in pink and yellow in panel B, respectively) of OHC from control heterozygous (black, gray) or homozygous *shaker2* (dark and light blue) littermates cultured in control conditions (darker points) or with tubocurarine (lighter points). Horizontal lines indicate the mean and statistical differences were obtained using Welch’s *t* test. For all panels, the age of explants is P5, and statistical significance is shown as: *P<0.05, **P<0.01, ***P<0.001, ****P<0.0001; n.s., not significant.

MET channel blockage also leads to the remodeling of the actin cytoskeleton in transducing stereocilia of IHC (Velez-Ortega et al., 2017). In heterozygous *shaker-2* IHC, SEM images showed thinning and shortening of stereocilia from the second row of the bundle, but no morphological changes were observed in stereocilia from the tallest row (Fig. 2A-C). In contrast, homozygous *shaker-2* IHC showed no evidence of cytoskeleton remodeling in stereocilia from the second (nor first) rows of the bundle. Homozygous *shaker-2* IHC stereocilia are not only abnormally short but they also lack a staircase arrangement (Stepanyan and Frolenkov, 2009, Hadi et al., 2020) thus the stereocilia height measurements from SEM images are rather challenging. Therefore, we also performed focused ion beam SEM (FIB-SEM) to obtain cross-sections of IHC stereocilia bundles and more accurate measurements of stereocilia heights from homozygous *shaker-2* IHC cells cultured in control conditions or with 60 µM tubocurarine for 5.5 hours (Fig. 2D). Once again, MET channel blockage did not lead to shortening of stereocilia from the first or second rows of the bundle.

**Figure 2.**
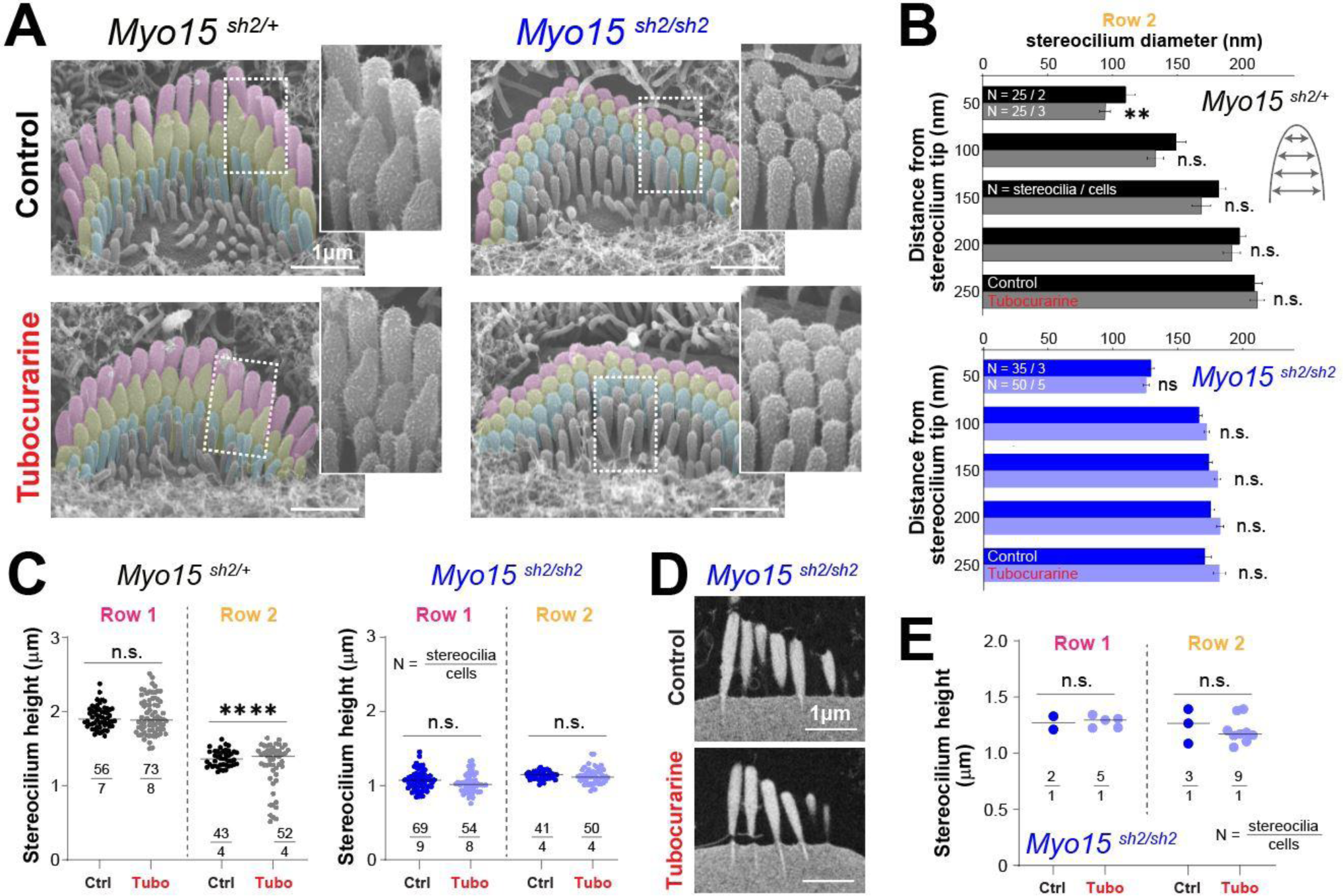
Inner hair cells lacking functional MYO15A do not exhibit mechanotransduction-dependent stereocilia remodeling. **(A)** Representative false-colored SEM images of inner hair cell (IHC) stereocilia bundles cultured for 4 hours at 37°C in control conditions (*top*) or with 300 µM tubocurarine (*bottom*), from heterozygous (*left*) or homozygous *shaker2* (*right*) littermates. Insets show highlighted bundle regions at higher magnification. **(B)** Diameter of stereocilia from the second row (colored in yellow in panel A) at several positions from the stereocilium tip (as indicated in cartoon), from IHC of control heterozygous (black, gray) or homozygous *shaker2* (dark and light blue) littermates cultured in control conditions (darker bars) or with tubocurarine (lighter bars). Data are shown as mean ± SE. Statistical differences were analyzed using a linear mixed model. **(C)** Heights of stereocilia from the first and second rows (colored in pink and yellow in panel A, respectively) of IHC from control heterozygous (black, gray) or homozygous *shaker2* (dark and light blue) littermates cultured in control conditions (darker points) or with tubocurarine (lighter points). Horizontal lines indicate the mean and statistical differences were obtained using Welch’s *t* tests. **(D)** Representative focused ion beam SEM (FIB-SEM) images from homozygous *shaker2* mice cultured at 37°C in control conditions (*top*) or with 60 µM tubocurarine (*bottom*) for 5.5 hours. **(E)** Heights of stereocilia from the first and second rows of the bundle obtained from FIB-SEM images of homozygous *shaker2* IHC cultured in control conditions (darker points) or with tubocurarine (lighter points). Horizontal lines indicate the mean and statistical differences were obtained using Welch’s *t* tests. For all panels, the age of explants is P5, and statistical significance is shown as: *P<0.05, **P<0.01, ***P<0.001, ****P<0.0001; n.s., not significant.

Our previous study (Velez-Ortega et al., 2017) showed quantitative larger stereocilia remodeling in OHC than in IHC with concentrations of blockers expected to reduce MET currents by ∼80-90% (*e.g*., 30 µM of tubocurarine). Here, using a 10X higher concentration of tubocurarine (300 µM, near saturating levels) we achieved remodeling in transducing stereocilia from IHC that was similar to that observed in OHC (compare Fig.1 to Fig. 2). This larger IHC remodeling, however, did not appear to be a pharmacological side effect triggered by higher tubocurarine concentrations since no evidence of stereocilia remodeling was observed in auditory hair cells from *Cib2* mutant mice lacking MET currents (Supp. Fig. 1 and 2).

To confirm the lack of MET-dependent stereocilia remodeling in the absence of functional MYO15A, we tested the effects of MET channel blockage on IHC and OHC from heterozygous and homozygous *shaker-2* mice after a longer incubation period (8 hours, Supp. Fig. 3A,B and 4A,B), a younger developmental age (P4, Supp. Fig. 3C,D and 4C,D) or a different MET channel blocker (30 M of benzamil for 5 hours, Supp. Fig. 3E,F and 4E,F). We also tested the effect of increased intracellular calcium buffering by culturing cells with the membrane-permeable calcium chelator BAPTA-AM (20 µM for 5 hours, Fig. 3E,F and 4E,F). Here and in our previous study, the effects of increased intracellular calcium buffering resembled those obtained with MET channel blockers (Velez-Ortega et al., 2017). In all these conditions, MET-dependent stereocilia remodeling was evident in transducing stereocilia from heterozygous but not homozygous *shaker-2* mice.

**Figure 3.**
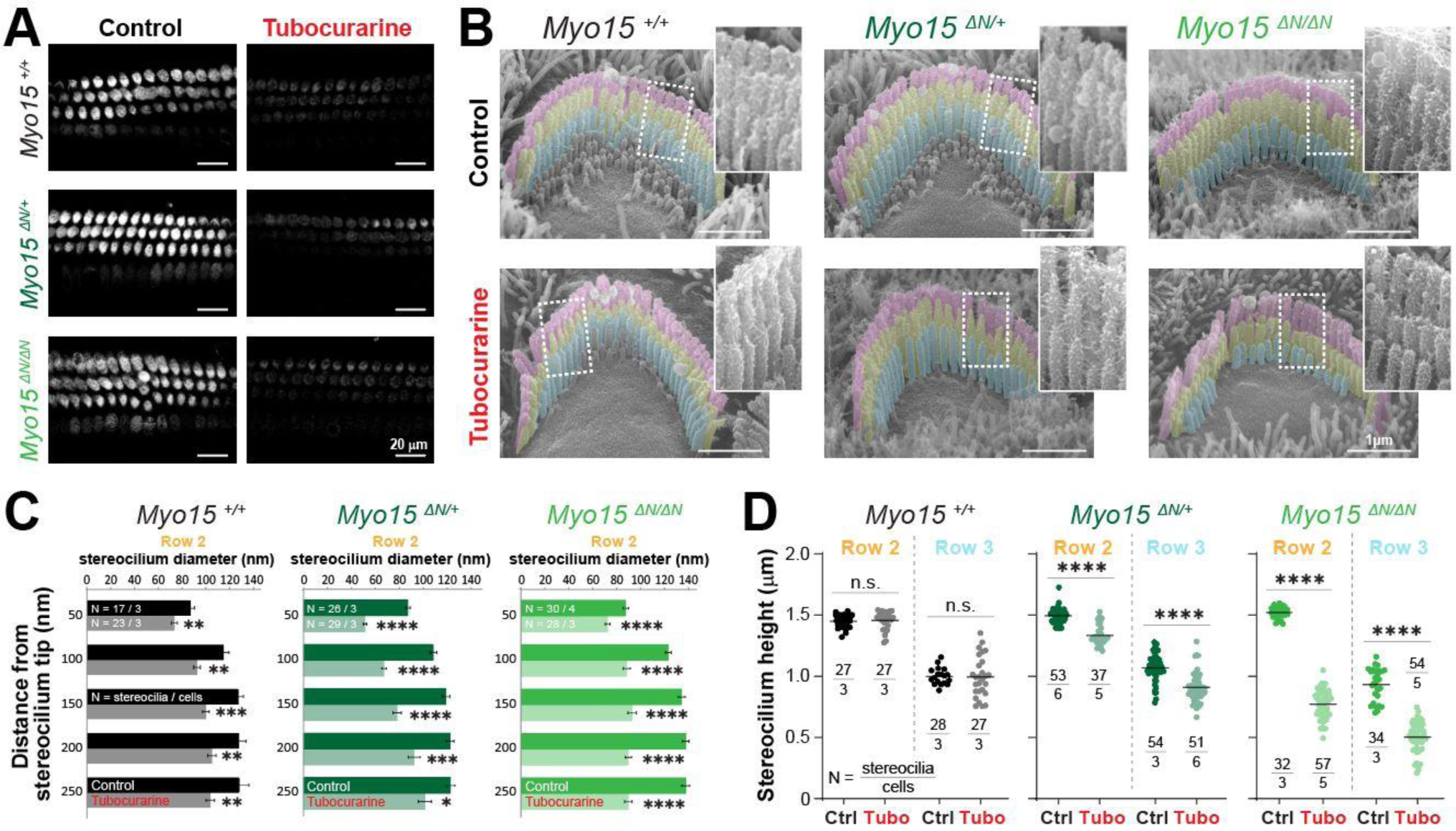
Outer hair cells lacking the long isoform of MYO15A show accelerated mechanotransduction-dependent stereocilia remodeling. **(A)** Decreased FM1-43 dye uptake in hair cells after application of the MET channel blocker, tubocurarine (300 µM), in wild-type *Myo15^+/+^* (*top*), heterozygous *Myo15^ΔN/+^* (*middle*) or homozygous *Myo15^ΔN/ΔN^* (*bottom*) littermates. **(B)** Representative false-colored SEM images of OHC stereocilia bundles cultured for 4 hours at 37°C in control conditions (*top*) or with 300 µM tubocurarine (*bottom*), from wild-type (*left*), heterozygous mice (*middle*) or homozygous *Myo15^ΔN/ΔN^* (*right*) littermates. Insets show highlighted bundle regions at higher magnification. **(C)** Diameter of stereocilia from the second row (colored in yellow in panel B) at several positions from the stereocilium tip from OHC of wild-type (black), heterozygous (darker shades of green) and homozygous *Myo15^ΔN/ΔN^* (brighter shades of green) littermates cultured in control conditions (darker bars) or with tubocurarine (lighter bars). Data are shown as mean ± SE. Statistical differences were analyzed using a linear mixed model. **(D)** Stereocilia heights from the second and third rows (colored in yellow and cyan, respectively, in panel B) of OHC from control wild-type (black, gray), heterozygous (darker shades of green) or homozygous *Myo15^ΔN/ΔN^* (brighter shades of green) littermates cultured in control conditions (darker points) or with tubocurarine (lighter points). Horizontal lines indicate the mean and statistical differences were obtained using a Šídák’s multiple comparisons test. For all panels, the age of explants is P7, and statistical significance is shown as: *P<0.01, **P<0.001, ***P<0.001, ****P<0.0001; n.s., not significant. Data are representative of 4 independent series with 3-4 hour incubations with 300 µM tubocurarine in P4-P7 explants.

We conclude that MYO15A is required for the delivery of the molecular machinery that regulates the remodeling of the stereocilia cytoskeleton in both IHC and OHC in response to changes in the influx of calcium through MET channels that are partially open at rest.

### 2.2 Transducing stereocilia exhibit exaggerated MET-dependent remodeling in the absence of the long isoform of MYO15A

Next, we sought out to explore whether the long isoform of MYO15A, which is typically enriched at the tips of transducing stereocilia (Fang et al., 2015), is involved in the MET-dependent remodeling of the stereocilia cytoskeleton. We used mice lacking the long isoform of MYO15A (*Myo15^ΔN/ΔN^*) which develop hair cell bundles with near normal staircase arrangements and MET currents (including resting currents) (Fang et al., 2015). These mice, however, exhibit progressive hearing loss and degeneration of stereocilia from the shorter rows in the bundle.

The entry of FM1-43 dye through MET channels open at rest was significantly blocked by tubocurarine in IHC and OHC of homozygous mice lacking the long isoform of MYO15A and their heterozygous and wild-type littermates (Fig. 3A), indicating the presence of resting MET currents in all three genotypes. We next exposed neonate organ of Corti explants to MET channel blockers and explored changes to the stereocilia morphology via SEM. In explants cultured in control conditions for 4 hours, mice lacking one (heterozygous *Myo15^ΔN/+^*) or two (homozygous *Myo15^ΔN/ΔN^*) alleles of the long isoform of MYO15A exhibited stereocilia bundles staircase arrangements similar to those observed in wild-type littermates (Fig. 3B and 4A). In OHC, the diameters of second row stereocilia were indistinguishable between all three genotypes (Supp. Fig. 5A) and so were the heights (comparisons against wild-type littermates: P = 0.1178 for heterozygous and P =0.9156 for homozygous mice). In IHC, however, second row stereocilia exhibited similar heights in all three genotypes (comparisons against wild-type littermates: P = 0.2774 for heterozygous and P = 0.2527 for homozygous mice) but the tip diameters were significantly smaller in mice lacking one or two alleles of the long isoform of MYO15A (Supp. Fig. 5B).

**Figure 4.**
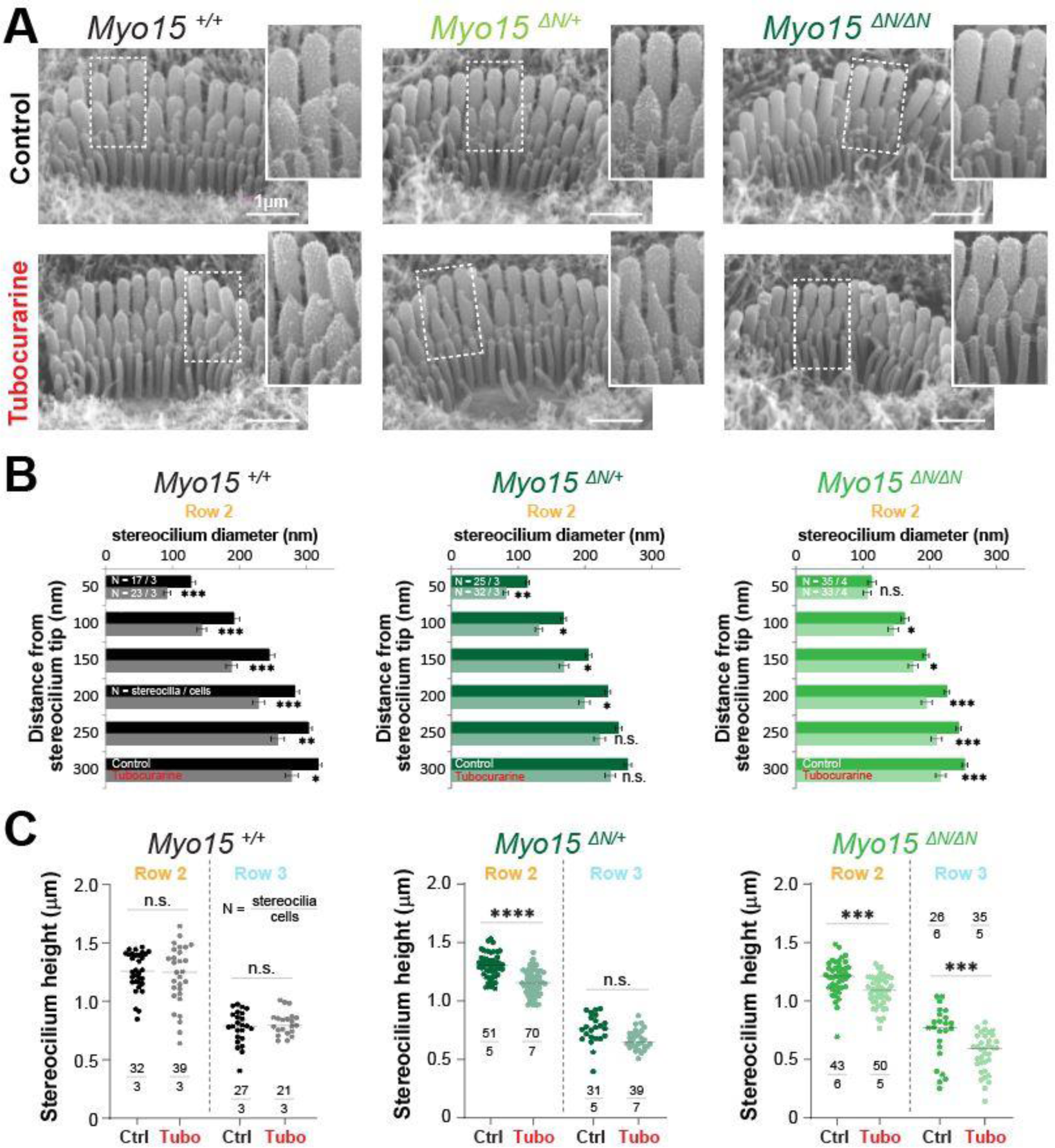
Inner hair cells lacking the long isoform of MYO15A show accelerated MET-dependent stereocilia remodeling. **(A)** Representative SEM images of IHC stereocilia bundles from wild-type (*left*), heterozygous (*middle*), or homozygous *Myo15^ΔN/ΔN^* (*right*) littermates cultured for 4 hours at 37°C in control conditions (*top*) or with 300 µM tubocurarine (*bottom*). Insets show highlighted bundle regions at higher magnification. **(B)** Diameter of stereocilia from the second row of IHC at several positions from the stereocilium tip, from wild-type (black), heterozygous (dark shades of green), and homozygous *Myo15^ΔN/ΔN^* (bright shades of green) littermates cultured in control conditions (darker bars) or with tubocurarine (lighter bars). Data are shown as mean ± SE. Statistical differences were analyzed using a linear mixed model. **(C)** Stereocilia heights from the second and third row of IHC from wild-type (black), heterozygous (darker shades of green), or homozygous *Myo15^ΔN/ΔN^*(brighter shades of green) littermates cultured in control conditions (darker points) or with tubocurarine (lighter points). Horizontal lines indicate the mean and statistical differences were obtained using Šídák’s multiple comparisons test. For all panels, the age of explants is P7, and statistical significance is shown as: *P<0.01, **P<0.001, ***P<0.001, ***P<0.0001; n.s., not significant. Data are representative of 4 independent series with 3-4 hour incubations with 300 µM tubocurarine in P4-P7 explants.

As expected in wild-type mice, a 4-hour incubation with the MET channel blocker tubocurarine (300 µM) led to significant thinning of the second row stereocilia in OHC and IHC bundles (Fig. 3B,C and 4A,B) but limited (often not yet significant) shortening of transducing second and third row stereocilia (Fig. 3D and 4C). In contrast, homozygous mice lacking the long isoform of MYO15A (*Myo15^ΔN/ΔN^*) showed significant shortening of transducing second and third row stereocilia, beyond the initial phase of stereocilia thinning (Fig. 3B-D and 4A-C). Mice lacking just one allele of the long isoform of MYO15A (*Myo15^ΔN/+^*) showed an intermediate response to MET channel blockage between wild-type and homozygous littermates (Supp. Fig. 5B,D) with significant thinning in the tips of second row stereocilia (Fig. 3B,C and 4A,B) as well as significant stereocilia shortening (Fig. 3D and 4C). These results were surprising given that mice lacking just one allele of the long isoform of MYO15A appear to have normal hearing (Fang et al., 2015).

Similar results were obtained after a longer incubation period (24 hours) with benzamil, a MET channel blocker pharmacologically unrelated to tubocurarine, at a 30 µM concentration which was shown to block ∼ 90% of MET currents induced by bundle deflections (Rusch et al., 1994). MET channel blockage induced significant shortening of transducing stereocilia in IHC and OHC from heterozygous and homozygous *Myo15^ΔN/ΔN^* mice and, once again, the degree of stereocilia shortening was significantly greater in the homozygous mice (Supp. Fig. 6).

Altogether, these results indicate that the long isoform of MYO15A delivers molecular machinery to the tips of transducing stereocilia crucial for proper cytoskeleton stability to avoid exaggerated MET-dependent stereocilia remodeling.

### 2.3 Stereocilia in the tallest row of the hair bundle exhibit MET-dependent remodeling when the long isoform of MYO15A is absent

The heights of stereocilia from the tallest row were shorter in mice lacking one or two alleles of the long isoform of MYO15A in OHC (Fig. 5A, comparisons against wild-type littermates: P = 0.02 for heterozygous and P < 0.0001 for homozygous mice) and IHC (Fig. 5C, comparisons against wild-type littermates: P = 0.04 for heterozygous and P = 0.008 for homozygous mice). In our previous study (Velez-Ortega et al., 2017), we showed that the MET-dependent remodeling of stereocilia was limited to the transducing rows of the bundle. Consistent with our previous study, no changes in stereocilia heights were observed in wild-type OHC or IHC cells after MET channel blockage with 300 µM of tubocurarine for 4 hours (Fig. 5A,C). To our surprise, careful examination of stereocilia bundle dimensions in our SEM images revealed significant shortening of stereocilia in the tallest row of IHC and OHC bundles in mice lacking one or both alleles of the long isoform of MYO15A (Fig. 5A,C). This MET-induced shortening of stereocilia from the tallest row of the bundle lacked the typical initial step of stereocilia tip thinning seen in transducing stereocilia (Fig. 3B and 4A). We did observe some small differences in the diameters of tallest row stereocilia from mice lacking one or both alleles of the long isoform of MYO15A, but these stereocilia tip diameters were largely maintained after MET channel blockage (Fig. 5B,D). MET channel blockage with 30 µM of benzamil for 24 hours also showed significant shortening of IHC and OHC stereocilia from the tallest row of the bundle in heterozygous and homozygous *Myo15^ΔN/ΔN^* littermates (Supp. Fig. 6).

**Figure 5.**
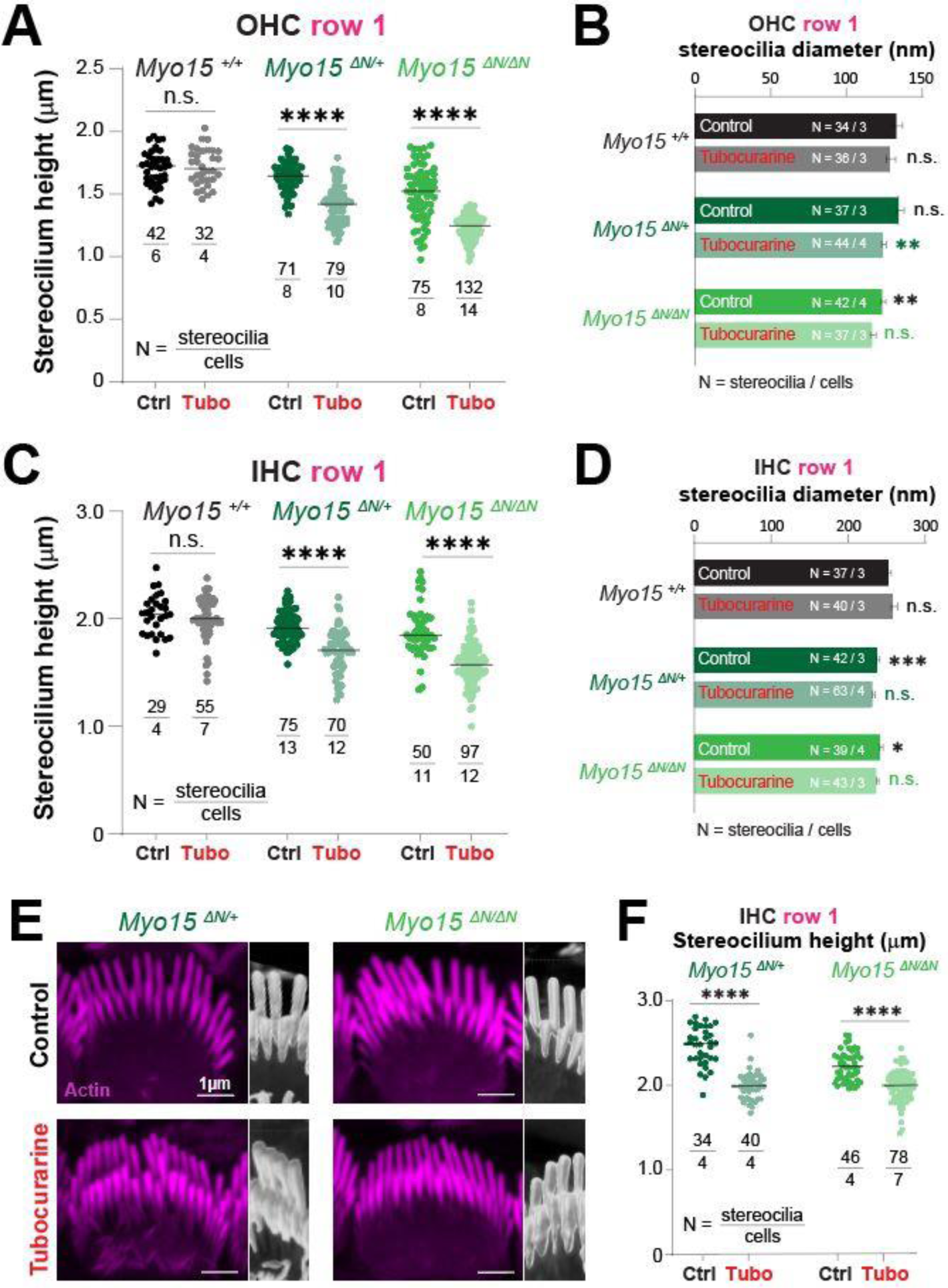
Hair cells lacking the long isoform of myosin XV exhibit MET-dependent stereocilia remodeling in the tallest row of the bundle. **(A-D)** Stereocilia heights **(A,C)** and diameters **(B,D)** from the first row of OHC **(A,B)** and IHC **(C,D)** from wild-type (black and gray), heterozygous (darker shades of green) or homozygous *Myo15^ΔN/ΔN^*(brighter shades of green) littermates that were cultured in control conditions (darker points/bars) or with 300 µM tubocurarine (lighter points/bars) for 4 hours at 37°C. Data are representative of 4 independent series with 3-4 hour incubations with 300 µM tubocurarine in P4-P7 explants. **(E)** Confocal images of IHC stereocilia stained against F-actin with fluorescently-labeled phalloidin (magenta), cultured for 4 hours at 37°C in control conditions (*top*) or with 300 µM tubocurarine (*bottom*), from heterozygous (*left*) and homozygous *Myo15^ΔN/ΔN^* (*right*) littermates. Insets show bundle regions with Imaris volume render at higher magnification. Images are representative of two independent experiments. **(F)** Stereocilia heights from the first row of IHC from heterozygous (dark shades of green) or homozygous *Myo15^ΔN/ΔN^* (bright shades of green) littermates cultured in control conditions (darker points) or with tubocurarine (lighter points). Stereocilia heights were measured from Imaris volume rendering of the confocal stacks. For all panels, the age of explants is P7, and statistical significance is shown as: *P<0.01, **P<0.001, ***P<0.001, ****P<0.0001; n.s., not significant. In panels **B** and **C**, significant differences are shown for control conditions between genotypes (black, relative to wild-type), or between control and tubocurarine conditions within each genotype (next to tubocurarine bars, color coded for each genotype: black for wild-type, dark green for heterozygous, and bright green for homozygous *Myo15^ΔN/ΔN^* littermates).

To confirm the shortening of stereocilia from the tallest row of the bundle that was triggered by MET channel blockage, we performed fluorescent labeling of F-actin and high-resolution confocal microscopy in mice lacking one or both alleles of the long isoform of MYO15 cultured with 300 M of tubocurarine for 4 hours. Volume rendering of the confocal stacks once again revealed a significant MET-induced decrease in the heights of stereocilia from the tallest row of IHC from heterozygous and homozygous mice lacking the long isoform of MYO15A (Fig. 5E,F).

We conclude that the absence of the long isoform of MYO15A results in decreased stability of the actin cytoskeleton within stereocilia from the tallest row of the bundle.

### 2.4 The absence of the long isoform of MYO15A leads to the mislocalization of several row identity proteins

Since the long isoform of MYO15A typically accumulates at the tips transducing stereocilia, we were surprised to observe MET-induced changes to the morphology of stereocilia in the tallest row of the bundle from mice lacking this long isoform. Therefore, we wondered whether the ‘row identity’ of stereocilia was affected. To test this, we performed immunolabeling against some known proteins that show preferential expression in the tips of stereocilia from either the tallest row or the transducing rows.

First, we evaluated the expression of ESPNL, a protein with high similarity to the crosslinker ESPN, predicted to have only one actin-binding domain, and detected during hair bundle development (up to ∼P11) as highly enriched in transducing stereocilia (Ebrahim et al., 2016). As expected, in wild-type IHC we observed labeling of ESPNL that was enriched in the transducing stereocilia at the two different developmental ages tested: P3 (Fig. 6A) and P7 (Fig. 6B). However, the ESPNL labeling in homozygous *Myo15^ΔN/ΔN^* IHC first appeared rather diffuse throughout the entire bundle (Fig. 6A) and was no longer detected by P7 (Fig. 6B). IHC from heterozygous *Myo15^ΔN/ΔN^* mice exhibited intermediate labeling patterns to those observed in wild-type and homozygous littermates (Fig. 6).

**Figure 6.**
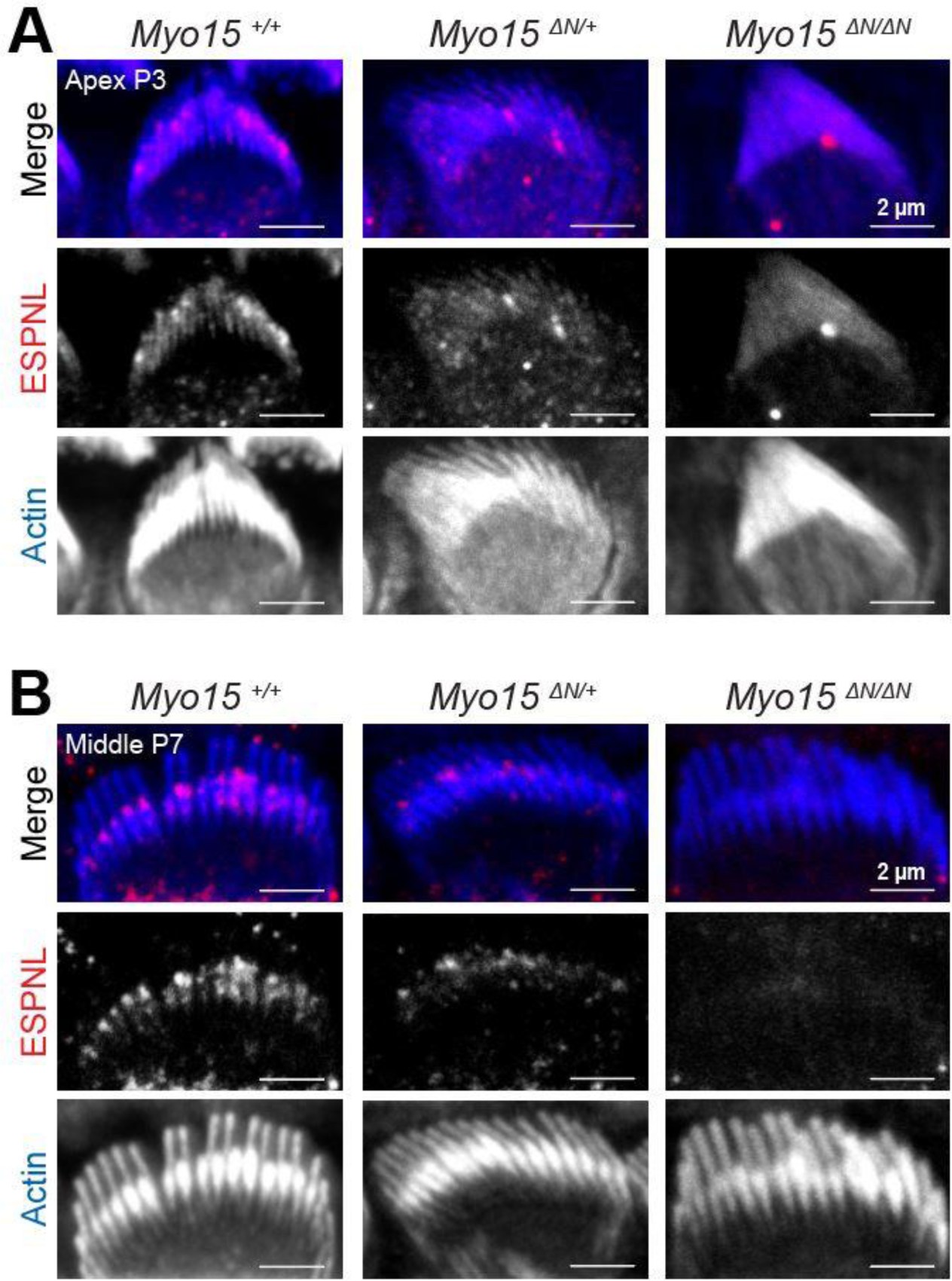
Decreased expression of ESPNL in inner hair cells lacking the long isoform of MYO15A. Maximum intensity projections of confocal stacks of IHC stereocilia from the apical **(A)** or middle **(B)** cochlear regions, at postnatal days 3 **(A)** or 7 **(B)**, from wild-type (*left*), heterozygous (*middle*), and homozygous *Myo15^ΔN/ΔN^* (*right*) mice immunolabeled against ESPNL (red) and counterstained against F-actin with fluorescently-labeled phalloidin (blue). Images are representative of two independent experiments, with similar labeling patterns along different cochlear regions.

Next, we evaluated whether proteins from the row 1 elongation complex were affected in mice lacking the long isoform of MYO15A. As described previously (Fang et al., 2015), the expression pattern of the actin capping/bundling protein EPS8 appears to be unaffected by the lack of the long isoform of MYO15A: it remains highly expressed at the tips of stereocilia from the tallest row in IHC and OHC from homozygous and heterozygous (*Myo15^ΔN/ΔN^*) mice, similar to what is observed in wild-type mice (Fig. 7A,B). At the time the *Myo15^ΔN/ΔN^* mice were first characterized, the role of the GPSM2-GNAI3 signaling complex in stereocilia elongation had not been reported. Thus, we next evaluated the expression pattern of GNAI3 in hair cells from mice lacking one or both alleles of the long isoform of MYO15A and in wild-type mice. While GNAI3 was still preferentially trafficked to the tips of stereocilia in the tallest row of the bundle, we noticed several stereocilia that lacked GNAI3 labeling in OHC from hetereozygous and homozygous *Myo15^ΔN/ΔN^* mice (Fig. 7C), while the GNAI3 labeling in IHC appeared normal (Fig. 7D). The GPSM2-GNAI3 signaling complex is also highly expressed in the bare zone of hair cells where it contributes to proper planar polarity (Tadenev et al., 2019), and we found similar GNAI3 labeling of the bare zone region in hair cells from mice lacking one or both alleles of the long isoform of MYO15A as in wild-type mice (Supp. Fig. 7).

**Figure 7.**
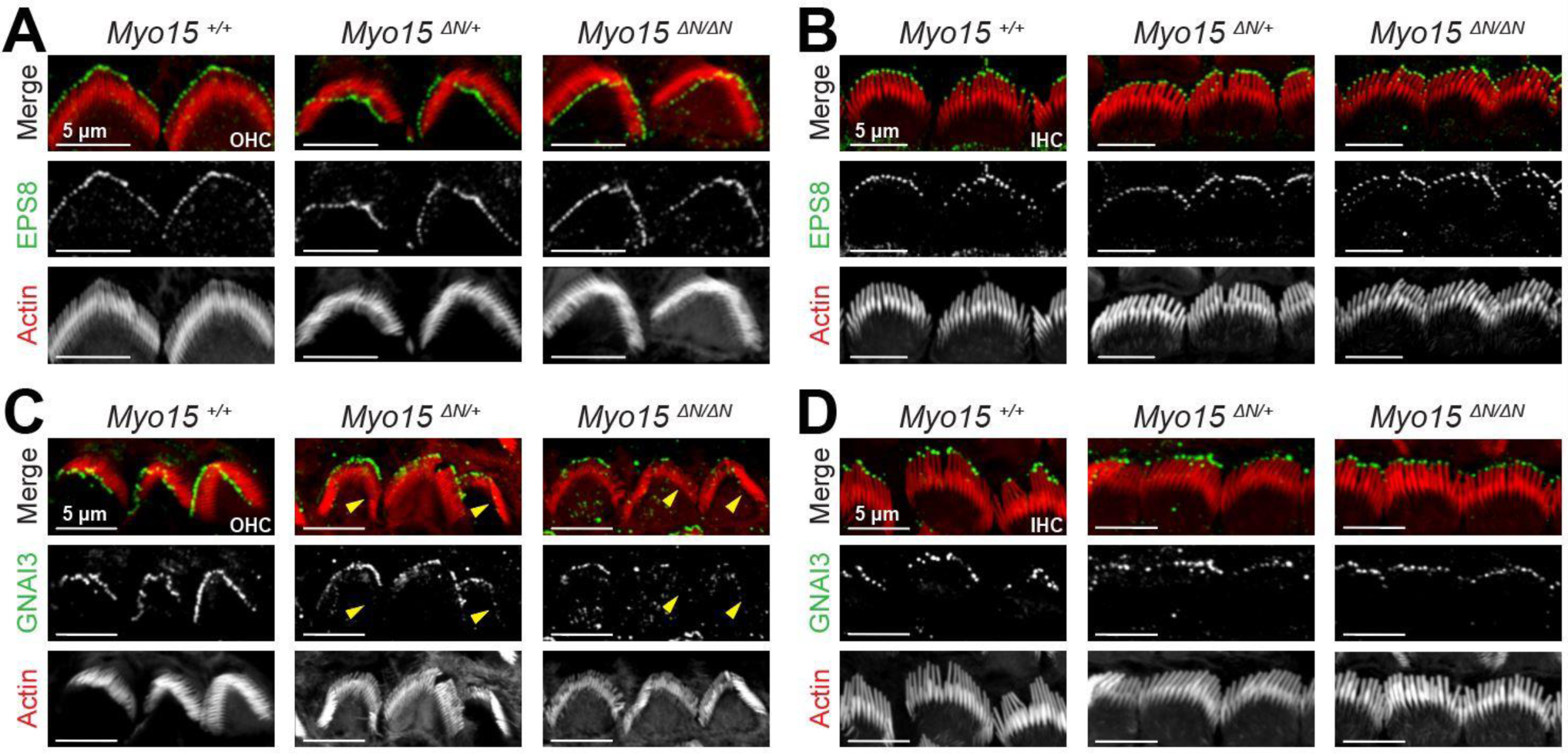
GNAI3 is mislocalized in outer, but not inner, hair cells lacking the long isoform of MYO15A. Maximum intensity projections of confocal stacks of OHC **(A,C)** and IHC **(B,D)** stereocilia from wild-type (*left*), heterozygous (*middle*), and homozygous *Myo15^ΔN/ΔN^*(*right*) mice immunolabeled against EPS8 (green) **(A,B)** or GNAI **(C,D)**, and counterstained against F-actin with fluorescently-labeled phalloidin (red). Yellow arrowheads point to OHC regions lacking GNA1 labeling in heterozygous and homozygous *Myo15^ΔN/ΔN^* mice. In panels **C** and **D**, the maximum intensity projections ignored Z planes near the cuticular plate to avoid confounding GNAI labeling from the bare zone of hair cells. To see the GNAI labeling in the bare zone, see Supplementary Figure 7. In all panels the age of explants is P7.

Our results indicate that, in the absence of the long isoform of MYO15A, several row identity proteins might not be trafficked properly.

## 3 Discussion

Our study is the first to show that MYO15A isoforms are necessary for the MET-dependent remodeling of the stereocilium actin cytoskeleton. We focused on the MET-driven plasticity of the stereocilium cytoskeleton of hair cells from the middle region of the mouse cochlea after postnatal day 4, once the initial stages of stereocilia elongation and thickening are completed and when row identity within the bundle has been clearly established. Under these conditions, the MET-dependent remodeling of the actin cytoskeleton is limited to the transducing shorter rows of stereocilia while stereocilia from the tallest row remains unaffected (Velez-Ortega et al., 2017, Krey et al., 2020). While no evidence of MET-driven remodeling was observed in hair bundles lacking all isoforms of MYO15A, the absence of the long isoform of MYO15A (isoform 1) led to exaggerated MET-dependent stereocilia remodeling in the shorter rows and to unexpected remodeling in the tallest row. These findings indicate that the ‘short’ or ‘middle’ isoforms of MYO15A (isoforms 2 and 3, respectively) deliver molecular machinery that enable stereocilia cytoskeleton rearrangements in response to changes in the resting MET current. However, at this point we cannot speculate on the specific MYO15A isoform that delivers such machinery because the expression levels and bundle localization of isoform 3 of MYO15A are still unknown. Nonetheless, this study highlights the need to explore whether isoform 3 of MYO15A is involved in the MET-driven remodeling of the stereocilium cytoskeleton using mice deficient in this isoform.

We did not observe any evidence of MET-dependent stereocilia remodeling in hair cells from *shaker-2* mice in which all isoforms of MYO15A are nonfunctional. Although we cannot completely rule out that remodeling could still happen in specific conditions not tested here, it is tempting to speculate that MYO15A isoforms either deliver (*i*) all molecular machinery required for the MET-driven remodeling of the stereocilium cytoskeleton, or (*ii*) machinery that works in series with proteins delivered by other myosin motors. In contrast, while the MET-dependent remodeling of stereocilia was present in the absence of the long isoform of MYO15A, this remodeling appeared to occur at a significantly faster rate. Therefore, the long isoform of MYO15A likely delivers molecular machinery that increases the cytoskeleton stability during the MET-driven remodeling of the stereocilium cytoskeleton. Since this isoform is preferentially trafficked to the shorter rows of the bundle, it was not surprising to observe changes in the degree of MET-dependent remodeling in the transducing stereocilia. In fact, shifting the balance towards greater MET-dependent stereocilium remodeling by decreasing the stability of the stereocilium cytoskeleton could also explain why mice deficient in the long isoform of MYO15A (*Myo15^ΔN/ΔN^*) exhibit premature stereocilia degeneration and subsequent hearing loss (Fang et al., 2015). Unexpectedly, hair cells from heterozygous *Myo15^+/ΔN^* mice –which develop normal hearing–, exhibited an intermediate MET-dependent stereocilia remodeling phenotype. Given the lack of a hearing phenotype in adult heterozygous *Myo15^+/ΔN^* mice, it is possible that the lack of one allele encoding the long isoform of MYO15A slightly alters the normal developmental stages of the hair bundle. In this case, the differences in MET-dependent stereocilium remodeling that we observed could simply be the result of comparisons between cells in slightly different developmental stages. Consistent with this hypothesis, we did observe significant differences in stereocilia diameters from the tallest and middle rows of IHC bundles and in the heights of IHC and OHC stereocilia from the tallest row between mice lacking one or two alleles of the long isoform of MYO15A and their wild-type littermates (Fig. 5D and Supp. Fig. 5C). Alternatively, adult heterozygous *Myo15^+/ΔN^* mice might have a hidden hearing phenotype. Perhaps the larger extent of MET-driven stereocilium remodeling works near a normal physiological range. However, under certain conditions –such as the noise-induced breakage of tip links–, the larger MET-dependent remodeling of transducing stereocilia could lead to some stereocilia degeneration. Therefore, it is worth exploring whether *Myo15^+/ΔN^* mice are more susceptible to noise-induced or age-related hearing loss than their wild-type littermates.

Unexpectedly, hair cells lacking one or both alleles of the long isoform of MYO15A also exhibited MET-dependent stereocilium remodeling in the tallest row of the bundle. MET-driven rearrangements of the actin cytoskeleton in stereocilia from the tallest row have been observed at early developmental bundle stages (*e.g*., at P4 in apical hair cells in (Krey et al., 2020)). Once again, these findings could indicate the presence of a delayed bundle development phenotype in the absence of the long isoform of MYO15A. However, if the bundle developmental stages are not affected, it is unlikely that the long isoform of MYO15A delivers molecular machinery that provides stability against MET-dependent remodeling to the tallest row, since this isoform is typically trafficked to the shorter rows of the bundle. Instead, it is plausible that the lack of expression of the long isoform of MYO15A affects the normal expression levels and/or distribution of isoform 3 of MYO15A within the bundle, which remains to be tested. A third hypothesis explaining the MET-dependent remodeling of stereocilia within the tallest row of the bundle in the absence of the long isoform of MYO15A is that, perhaps, some MET channels are mislocalized to the upper end of the tip link. In this scenario, stereocilia from the tallest row would also experience a resting MET current. Since the MET-driven shortening of stereocilia in the tallest row is rather uniform, it would indicate that all stereocilia within the tallest row have MET channels at the upper tip link end. Thus, could hair cells lacking the long isoform of MYO15A possibly have larger MET currents upon bundle deflections? Previously, abnormal current-displacement MET curves were described for *Myo15^ΔN/ΔN^* mice relative to their *Myo15^+/ΔN^* littermates (Fang et al., 2015). Larger maximum MET currents were indeed reported for OHC (and perhaps IHC at larger displacements) in *Myo15^+/ΔN^* mice. However, that study did not evaluate wild-type littermates. In addition, MET currents were assessed with a stiff probe, which did not allow for proper measurements of resting MET currents. Therefore, the presence of abnormal resting MET currents cannot be ruled out in mice lacking one or two alleles of the long isoform of MYO15A.

An additional interesting observation regarding the MET-dependent remodeling of stereocilia from the tallest row was the lack of stereocilia thinning at the tips, like we typically observe in stereocilia from the shorter rows of the bundle. In the presence of tip links, the MET-dependent remodeling of transducing stereocilia starts with significant thinning of stereocilia tips and eventual shortening. Previously, we only observed uniform shortening of transducing stereocilia (not preceded by stereocilium tip thinning) after tip link breakage with BAPTA-buffered extracellular medium (Velez-Ortega et al., 2017). We hypothesized that the thinning at the stereocilium tip was due to the partial blockage (∼70-90%) of MET channels. We thought the small influx of calcium ions was able to stabilize actin filaments near the MET channel while actin filaments at the periphery of the stereocilium –far from the MET channel and experiencing low intracellular calcium concentrations– would depolymerize first. Here, we used a saturating concentration of MET channel blocker, yet we still observed significant thinning at the stereocilium tips instead of uniform stereocilium shortening. These results likely indicate the presence of additional signaling pathways provided by the tip links that help stabilize the stereocilium actin cytoskeleton. In support of this hypothesis, a recent study shows that tip-link mutant mice exhibit some differences in stereocilia bundle dimensions that cannot solely be explained by defects in MET currents (Krey et al., 2023).

The exact molecular machinery delivered by MYO15A isoforms to allow and fine-tune the extent of MET-dependent remodeling in the stereocilium cytoskeleton remains to be fully explored. We did observe a decrease in the expression of ESPNL in transducing stereocilia from mice lacking one or two alleles of the long isoform of MYO15A. Although ESPNL is a known cargo protein of MYO3A (Ebrahim et al., 2016), its expression is still present at the tips of stereocilia from hair cells lacking both MYO3A and MYO3B (Lelli et al., 2016). However, it remains to be explored whether ESPNL could be a cargo of MYO15A. Given the increased levels of MET-dependent remodeling in transducing stereocilia from mice lacking the long isoform of MYO15A, perhaps ESPNL is required to increase the stability of the stereocilium actin cytoskeleton. In support of this hypothesis, mice lacking ESPNL show degeneration of transducing stereocilia (Ebrahim et al., 2016).

Due to the unexpected MET-driven remodeling observed in stereocilia from the tallest row of the hair bundle in mace lacking one or two alleles of the long isoform of MYO15A, we also explored the expression levels of GNA13, a ‘row-identity’ signaling protein trafficked to the tips of stereocilia from the tallest row. GNAI3 is detected in some stereocilia tips by E18.5 and in virtually all stereocilia tips by P4 (Tadenev et al., 2019). In *Myo15^+/ΔN^* and *Myo15^ΔN/ΔN^* mice, however, GNAI3 labeling was still lacking in several OHC stereocilia by P7. Once again, these results could be consistent with defects in the proper stages of bundle development and the establishment of stereocilium row identity within the bundle of mice lacking one or two alleles of the long isoform of MYO15A. In addition, the GPSM2-GNAI3 signaling complex might be crucial for the normal stability of the actin cytoskeleton in stereocilia from the tallest row of the bundle and its insensitivity to changes in resting MET current levels.

Altogether, our results indicate that MYO15A isoforms deliver to the tips of shorter stereocilia molecular machinery required to allow the remodeling of the actin cytoskeleton in response to changes in resting MET currents. In addition, the long isoform of MYO15A also delivers machinery to increase cytoskeleton stability –thus preventing exaggerated MET-dependent stereocilium remodeling– and influences the row identity of stereocilia.

## 4 Methods

### 4.1 Mice and cochlear explants

Organ of Corti explants from *Myo15^sh2/sh2^*, *Myo15^ΔN/ΔN^, CIB2^F91S/F91S^* and *CIB2^tm1a/tm1a^* mice along with their heterozygous and wild-type littermates were isolated at postnatal days 4 through 7. *Myo15^sh2/sh2^* mice are maintained in the laboratory in an inbred mixed background (B6N-Tyr^c-Brd^/BrdCrCrl and C57Bl/6J). *Myo15^ΔN/ΔN^* mice generated by Dr. Jonathan Bird (REF) are maintained in a B57Bl/6 background. *CIB2^F91S/F91S^* and *CIB2^tm1a/tm1a^* mice generated by Dr. Zubair Ahmed (REF) and maintained in a B57Bl/6J background were kindly provided to us by Dr. Gregory Frolenkov. Temporal bones from the mice were isolated, and access to the cochlea was gained by removing surrounding bone. Following the bone removal, the modiolus and stria vascularis were subsequently removed. The cochlear explants were held in place during incubations and live-cell imaging by two flexible glass fibers glued to the bottom of a plastic Petri dish.

The University of Kentucky Institutional Animal Care and Use Committee granted approval for all animal procedures (protocol 2020-3535 to A.C.V.)

### 4.2 Incubations with MET channel pharmacological blockers

Organ of Corti explants were cultured in DMEM (Gibco, 11-995-065) supplemented with 7% FBS (Gibco, 16140071) and 10 mg/mL ampicillin (Calbiochem, 171254), at 37°C and 5% CO_2_ for up to 24 hours with 30 µM of benzamil (Sigma, B2417) in DMSO (VWR, N182-5X), with 60 or 300 µM of tubocurarine (Sigma Aldrich, 93750), 20 µM of BAPTA-AM (Invitrogen, B6769) pre-mixed with 20% Pluronic F-127 solution in DMSO (Invitrogen, P3000MP), or in vehicle-control conditions (0.05% DMSO). All samples had their tectorial membrane removed after incubation and were fixed in either 4% paraformaldehyde (PFA) (for subsequent immunolabeling), formaldehyde/glutaraldehyde (3% each) in 0.1 M sodium cacodylate buffer (Electron Microscopy Sciences, 15950) supplemented with 2 mM of CaCl_2_ (Sigma-Aldrich, 16538-06) (for scanning electron microscopy) or 2% paraformaldehyde/2.5% glutaraldehyde in 0.1 M sodium cacodylate buffer (Electron Microscopy Sciences, 15960-01) with 1% tannic acid (Sigma Aldrich, 403040) (for focused ion beam electron microscopy).

### 4.3 FM1-43 dye uptake

Freshly-isolated organ of Corti explants were incubated for 30 seconds with cold Ca^2+^/Mg^2+^-free Hank’s balanced salt solution (HBSS) (Gibco, 14175-079) supplemented with 5 µM of FM1-43 dye (ThermoFisher, T35356) in the absence or presence of the MET channel blocker tubocurarine. The explants were rinsed several times with cold Leibovitz’s L-15 (Gibco, 21-083-027) and imaged immediately. Images were acquired using with a Leica SP8 upright confocal microscope equipped with a Leica XCX APO L 63X 0.9 NA water immersion objective.

### 4.4 Immunolabeling and confocal microscopy

All samples were fixed with cold 4% PFA supplemented with 20 mM CaCl_2_.samples were left in the fixative solution at least 24 hours at 4°C. Samples then were rinsed in PBS (Gibco, 10010023) and then permeabilized in 0.5% Triton™ X-100 (Electron Microscopy Sciences, 22142) in PBS for 1 hour. Samples were next blocked for two hours with a solution of 5% normal goat serum (Invitrogen,10000c), 2% bovine serum albumin (BSA, Thermofisher scientific, 37525) and 0.25% Triton™ X-100. Samples were incubated overnight with primary antibodies against EPS8 (BD Biosciences, 610143, at 1:100), GNAI3 (Millipore Sigma, G4040), or ESPNL (BG35961 at 1:50, previously characterized in (Ebrahim et al., 2016)) at 4°C. Samples were rinsed with PBS and 0.2% BSA and incubated with fluorescently labeled secondary antibodies (Alexa Fluor 555 goat anti-mouse, ThermoFisher Scientific, A-21127, at 1:100; CF 555 goat anti-rabbit, Sigma-Aldrich, SAB4600068, at 1:100; or Alexa Fluor 488 goat anti-rabbit, ThermoFisher Scientific, A-11034, at 1:1000) for three hours at 120 rpm at room temperature. The samples were counterstained at room temperature with either rhodamine phalloidin (1 unit, ThermoFisher Scientific, R415) for 30 minutes or Alexa Fluor 488 phalloidin (1 unit, ThermoFisher Scientific, A12379) for 45 minutes. Tissues were rinsed in PBS, mounted in ProLong Diamond antifade (ThermoFisher Scientific, P36965) and allowed to cure for 72 hours before imaging. Images were acquired from the middle cochlear region with a Leica SP8 upright confocal microscope equipped with a Leica HCX PL APO 100X 1.44 NA objective lens. The typical voxel size used during imaging acquisition of hair cell bundles was ∼20 nm in X and Y, and 50 nm in Z. Imaging drifts were corrected in Huygens software.

### 4.5 Scanning electron microscopy (SEM)

Organ of Corti explants were fixed in cold formaldehyde/glutaraldehyde (3% each) in 0.1 M sodium cacodylate buffer supplemented with 2 mM of CaCl_2_ as described above. Samples were kept in fixative at 4°C for at least 24 hours until further processing. Distilled water was used to rinse the samples, followed by dehydration through a graded series of ethanol (VWR, 89125-170). Subsequently, samples were subjected to critical point drying from liquid CO_2_ (Leica, CPD300) and coated with 5 nm platinum through sputter coating (Quorum Technologies, Q150T). Hair cells from the middle cochlear region (near the 50% point in cochlear length) were imaged with a field-emission scanning electron microscope (FEI, Helios Nanolab 660). This work was performed in part at the U.K. Electron Microscopy Center, a member of the National Nanotechnology Coordinated Infrastructure (NNCI), which is supported by the National Science Foundation (NNCI-2025075).

For accurate stereocilia height measuring, images of the same bundle were obtained from different angles, including views from the lateral (back) and medial (front) sides of the bundle. Measurements were performed on Fiji ImageJ. Calculations of the heights of each stereocilium were performed as described in (Velez-Ortega et al., 2017).

### 4.6 Focused ion beam scanning electron microscopy (FIB-SEM)

Organ of Corti explants were fixed in 2% paraformaldehyde/ 2.5% glutaraldehyde (Electron Microscopy Sciences, 15960-01) supplemented in 1% tannic acid (Sigma Aldrich, 403040). Distilled water was used to rinse the samples, followed by a graded series of glycerol (Electron Microscopy Sciences, 16550). Samples underwent plunge freezing using Freon, and liquid nitrogen. Samples were then moved to a 1:1 solution of liquid nitrogen and 1% uranium acetate (Electron Microscopy Sciences, 22400) before undergoing freeze substitution and low temperature embedding (Leica, EMAFS2) with graded series of methanol (Electron Microscopy Sciences, 18511), lowicryl (Electron Microscopy Sciences, 14330) and curing with UV light. Resin blocks were thinned (Leica, SM2000R), trimmed (Boeckeler Instruments, PTXLPowerTome) and coated with 25 nm platinum through sputter coating (Quorum Technologies, Q150T). Samples were then milled with a focused ion beam (65 nA) at 20 nm steps and imaged (near the mid-point of the cochlea) with a field-emission scanning electron microscope (FEI, Helios Nanolab 660) at the U.K. Electron Microscopy Center.

### 4.7 Statistical analysis

Data is shown as Mean ± SEM, unless otherwise noted. Asterisks denote statistical significance from either Welch’s *t* tests, Mann-Whitney tests, Šídák’s multiple comparisons tests, or linear mixed effect models using Prism (GraphPad Software) or Origin (OriginLab Corp.). Results are shown with P values < 0.05 (*), < 0.01 (**), < 0.001 (***) or < 0.0001 (****) and the specific test is indicated in each figure legend.

## 6 Author Contributions

A.C.V. and A.I.L. conceived and designed the study, and wrote the manuscript. A.I.L. performed organ of Corti treatments, preparation of samples for electron microscopy, electron microscopy imaging, immunostaining, mechanotransduction assessment using FM1-43 dye, confocal microscopy imaging, and all data analysis. A.M.K. performed immunostaining and confocal microscopy imaging. M.T.M. performed focused-ion beam-scanning electron microscopy sample preparation and imaging.

## 8 Acknowledgments

The authors would like to thank Dr. Peter Barr-Gillespie for providing ESPNL antibodies; Drs. Zubair Ahmed and Gregory Frolenkov for providing the mechanotransduction-deficient (*Cib2* mutant) mice; Dr. Basile Tarchini for suggestions on antibodies against GNAI3 and GPSM2; and Drs. Jonathan Bird and Gregory Frolenkov for helpful discussions and comments throughout the execution of this study.

## 9 Data Availability Statement

All data generated in this study is available from the corresponding author upon request.

