## Supplementary Figures for "Myosin XVA isoforms participate in the mechanotransduction-dependent remodeling of the actin cytoskeleton in auditory stereocilia"

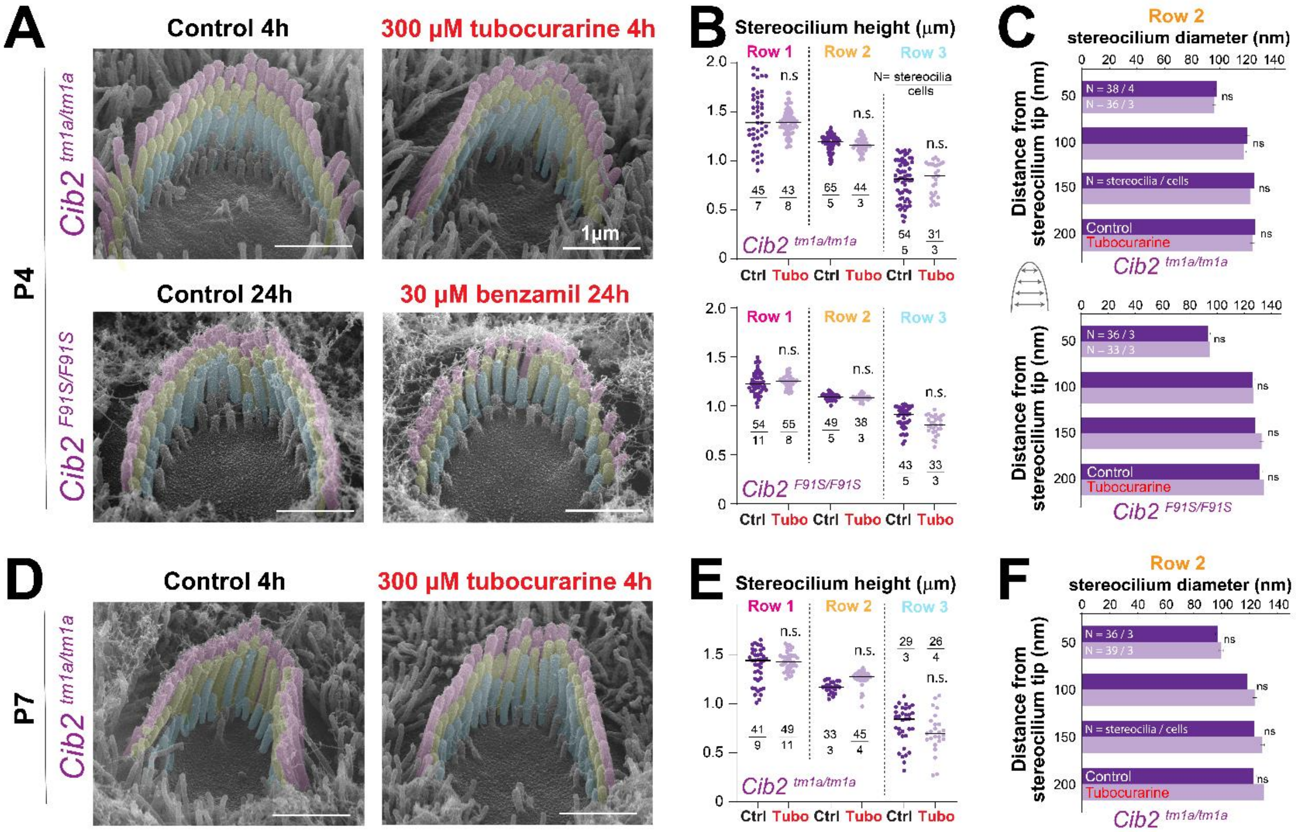


**Supplementary Figure 1. Stereocilia cytoskeleton remodeling in OHC is not a side effect of the pharmacological MET channel blockers**. **(A,D)** Representative false-colored SEM images of OHC bundles from mechanotransduction-deficient mice *Cib2*$^{tm1a/tm1a}$ (**A** *top*, **D**) and *Cib2*$^{F91S/F91S}$ (**A** *bottom*) cultured in control conditions (*left*) or in the presence of MET channel blockers (*right*): 300 µM tubocurarine (**A** *top*, **D**) or 30 µM benzamil (**A** *bottom*) for 4 (**A** *top*, **D**) or 24 hours (**A** *bottom*). **(B,E)** Heights of OHC stereocilia from the first, second, and third rows (colored in pink, yellow, and cyan, respectively, in panels **A** and **D**) from mechanotransduction-deficient mice *Cib2*$^{tm1a/tm1a}$ (**B** *top*, **E**) and *Cib2*$^{F91S/F91S}$ (**B** *bottom*) cultured in control conditions (dark purple points) or in the presence of MET channel blockers (light purple points): 300 µM tubocurarine (**B** *top*, **E**) or 30 µM benzamil (**B** *bottom*) for 4 (**B** *top*, **E**) or 24 hours (**B** *bottom*). Horizontal lines indicate the mean. **(C,F)** Diameters of OHC stereocilia from the second row (colored in yellow in panels **A** and **D**) from mechanotransduction-deficient mice *Cib2*$^{tm1a/tm1a}$ (**C** *top*, **F**) and *Cib2*$^{F91S/F91S}$ (**C** *bottom*) cultured in control conditions (dark purple bars) or in the presence of MET channel blockers (light purple bars): 300 µM tubocurarine (**C** *top*, **F**) or 30 µM benzamil (**C** *bottom*) for 4 (**C** *top*, **F**) or 24 hours (**C** *bottom*). Data are shown as mean ± SE. The age of explants is P4 (**A-C**) and P7 (**D-F**). Statistical differences were obtained using Welch's *t* tests; n.s., non-significant.

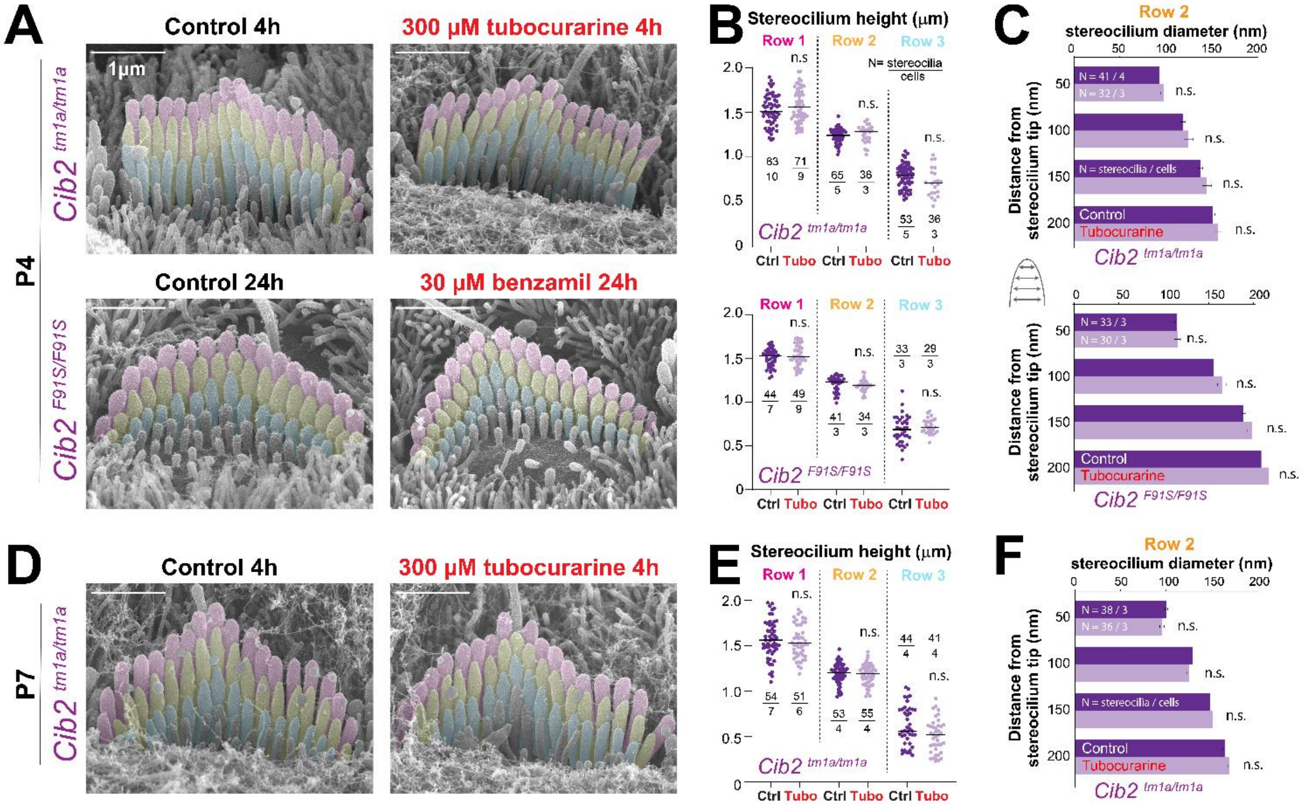


**Supplementary Figure 2. Stereocilia cytoskeleton remodeling in IHC is not a side effect of the pharmacological MET channel blockers**. **(A,D)** Representative false-colored SEM images of IHC bundles from mechanotransduction-deficient mice $Cib2^{tm1a/tm1a}$ (**A** *top*, **D**) and $Cib2^{F91S/F91S}$ (**A** *bottom*) cultured in control conditions (*left*) or in the presence of MET channel blockers (*right*): 300 µM tubocurarine (**A** *top*, **D**) or 30 µM benzamil (**A** *bottom*) for 4 (**A** *top*, **D**) or 24 hours (**A** *bottom*). **(B,E)** Heights of IHC stereocilia from the first, second, and third rows (colored in pink, yellow, and cyan, respectively, in panels **A** and **D**) from mechanotransduction-deficient mice $Cib2^{tm1a/tm1a}$ (**B** *top*, **E**) and $Cib2^{F91S/F91S}$ (**B** *bottom*) cultured in control conditions (dark purple points) or in the presence of MET channel blockers (light purple points): 300 µM tubocurarine (**B** *top*, **E**) or 30 µM benzamil (**B** *bottom*) for 4 (**B** *top*, **E**) or 24 hours (**B** *bottom*). Horizontal lines indicate the mean. **(C,F)** Diameters of IHC stereocilia from the second row (colored in yellow in panels **A** and **D**) from mechanotransduction-deficient mice $Cib2^{tm1a/tm1a}$ (**C** *top*, **F**) and $Cib2^{F91S/F91S}$ (**C** *bottom*) cultured in control conditions (dark purple points) or in the presence of MET channel blockers (light purple points): 300 µM tubocurarine (**C** *top*, **F**) or 30 µM benzamil (**C** *bottom*) for 4 (**C** *top*, **F**) or 24 hours (**C** *bottom*). Data are shown as mean ± SE. The age of explants is P4 (**A-C**) and P7 (**D-F**). Statistical differences were obtained using Welch's *t* tests; n.s., non-significant.

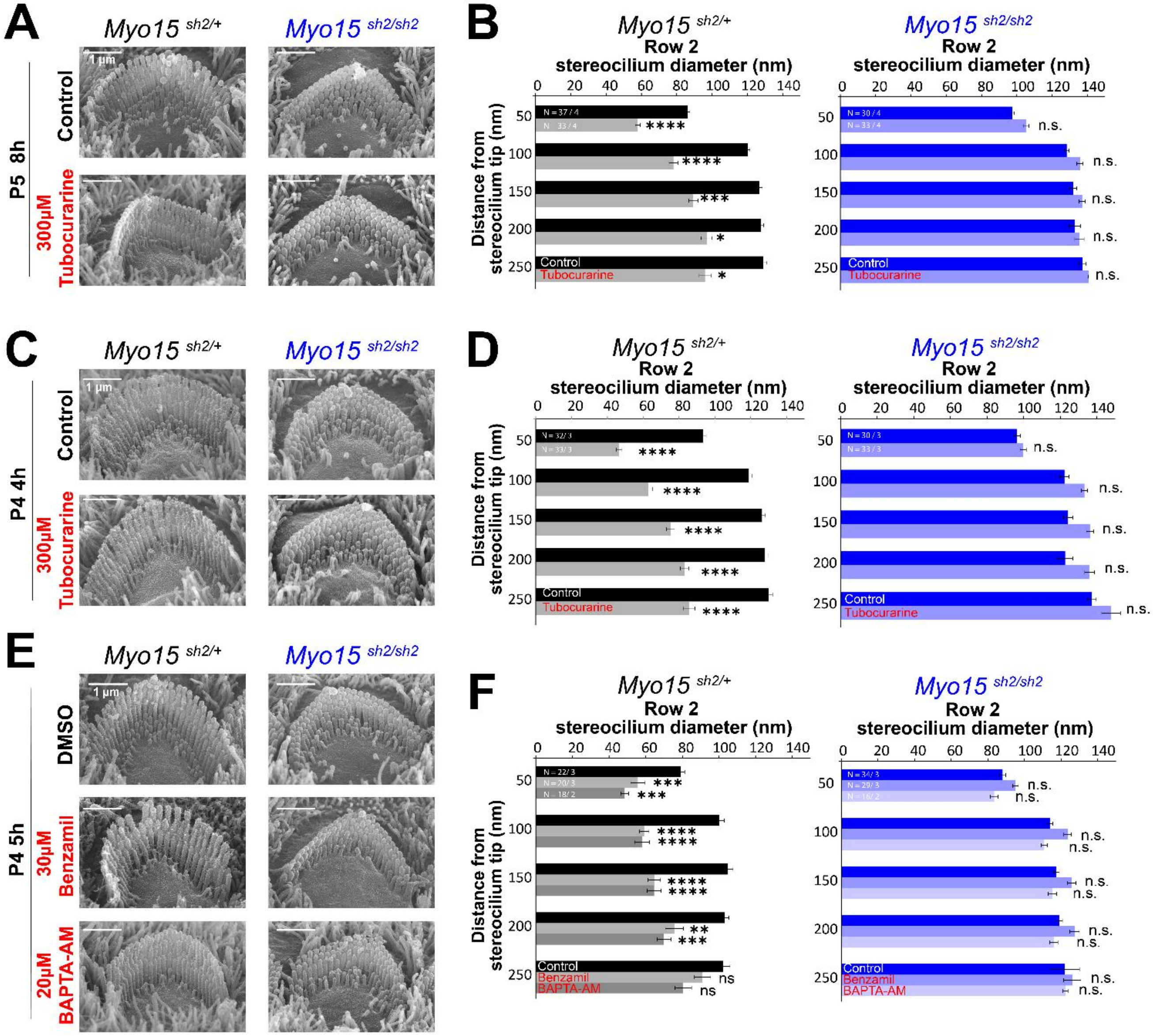


**Supplementary Figure 3. Lack of MET-dependent stereocilia remodeling in OHC from mice lacking functional MYO15A. (A,C,E)** Representative SEM images of OHC from heterozygous (*left*) or homozygous *shaker-2* (*right*) mice cultured in control conditions (*top* row in each panel) or in the presence of 300 µM tubocurarine for 8 (**A** *bottom*) or 4 (**C** *bottom*) hours, 30 µM benzamil for 5 hours (**E** *middle*), or 20 µM BAPTA-AM for 5 hours (**E** *bottom*). **(B,D,F)** Diameters of OHC stereocilia from the second row from heterozygous (*left*, black and gray) or homozygous *shaker-2* (*right*, shades of blue) mice cultured in control conditions (dark bars) or in the presence of 300 µM tubocurarine for 8 (**B,** gray and light blue) or 4 (**D,** gray and light blue) hours, 30 µM benzamil for 5 hours (**F**, light gray and medium shade of blue), or 20 µM BAPTA-AM for 5 hours (**F,** dark gray and lightest shade of blue). Data are shown as mean ± SE. The age of explants is P5 (**A,B**) and P4 (**C-F**). Statistical differences were obtained using a linear mix model analysis, and are shown as: *P<0.05, **P<0.01, ***P<0.001, ****P<0.0001; n.s., not significant.

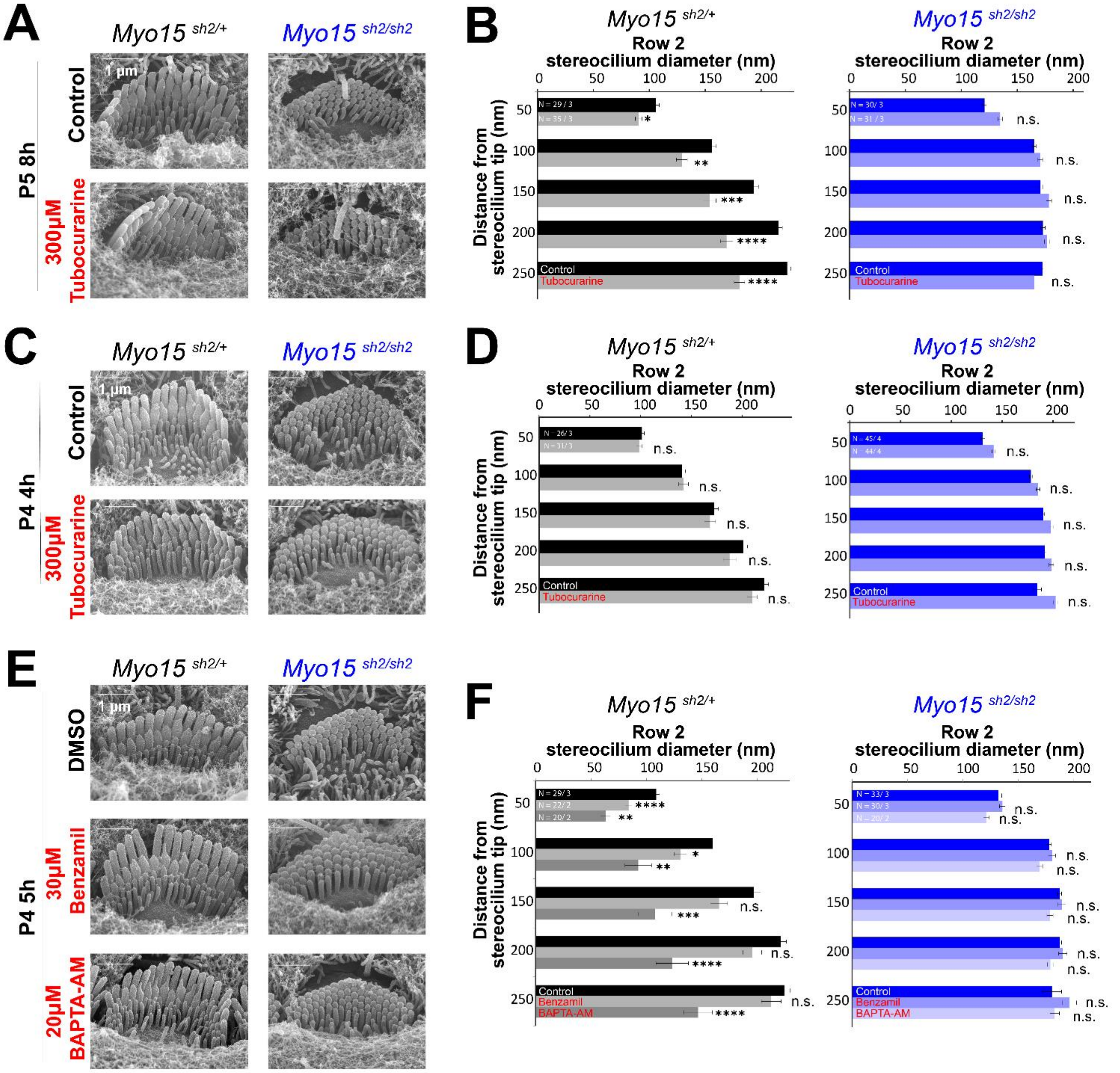


**Supplementary Figure 4. Lack of MET-dependent stereocilia remodeling in IHC from mice lacking functional MYO15A. (A,C,E)** Representative SEM images of IHC from heterozygous (*left*) or homozygous *shaker-2* (*right*) mice cultured in control conditions (*top* row in each panel) or in the presence of 300 µM tubocurarine for 8 (**A** *bottom*) or 4 (**C** *bottom*) hours, 30 µM benzamil for 5 hours (**E** *middle*), or 20 µM BAPTA-AM for 5 hours (**E** *bottom*). **(B,D,F)** Diameters of IHC stereocilia from the second row from heterozygous (*left*, black and gray) or homozygous *shaker-2* (*right*, shades of blue) mice cultured in control conditions (dark bars) or in the presence of 300 µM tubocurarine for 8 (**B,** gray and light blue) or 4 (**D,** gray and light blue) hours, 30 µM benzamil for 5 hours (**F**, light gray and medium shade of blue), or 20 µM BAPTA-AM for 5 hours (**F,** dark gray and lightest shade of blue). Data are shown as mean ± SE. The age of explants is P5 (**A,B**) and P4 (**C-F**). Statistical differences were obtained using a linear mix model analysis, and are shown as: *P<0.05, **P<0.01, ***P<0.001, ****P<0.0001; n.s., not significant.

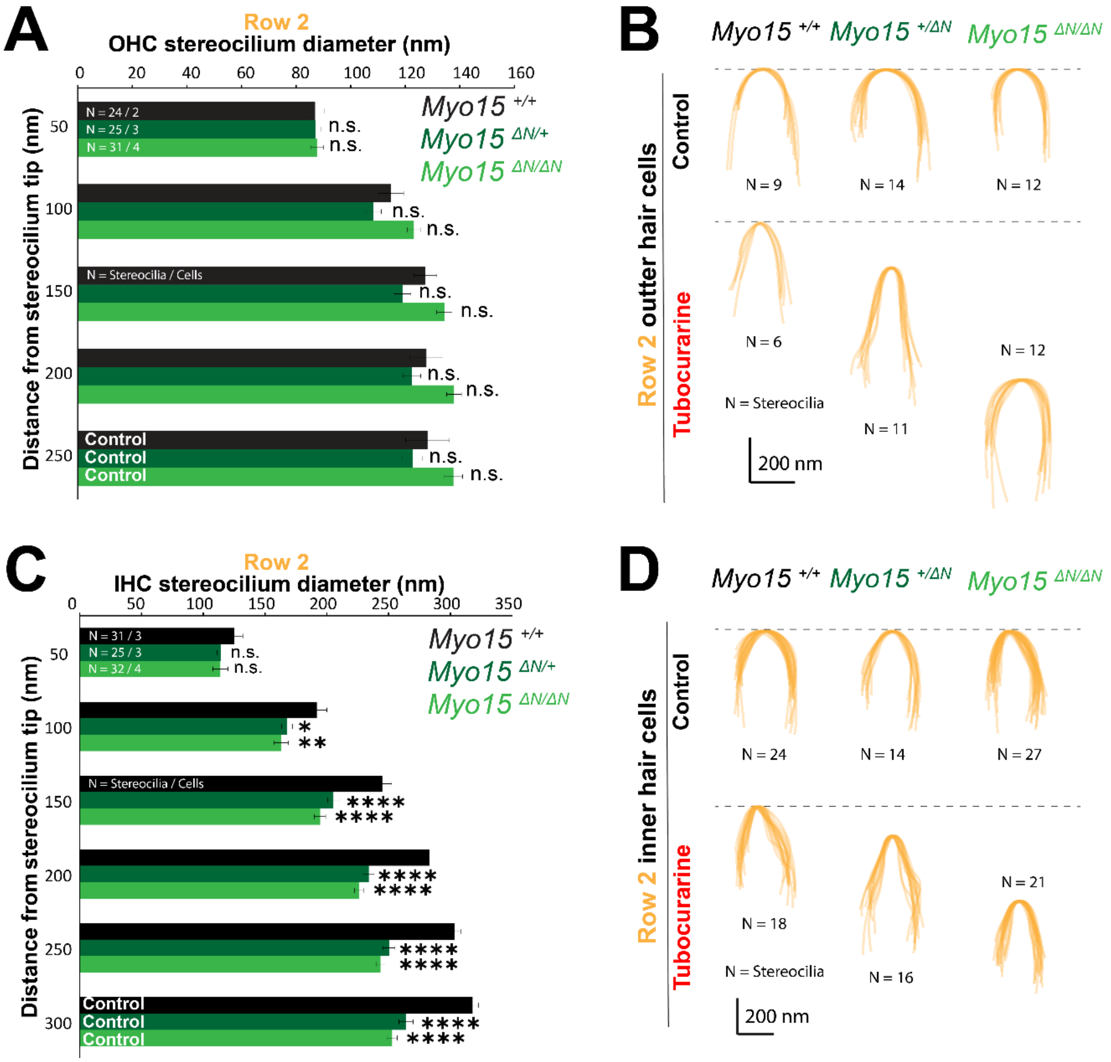


**Supplementary Figure 5. The long isoform of MYO15A confers stability to the second row stereocilia cytoskeleton**. **(A,C)** Diameter of stereocilia from the second row of OHC **(A)** and IHC **(C)** at several positions from the stereocilium tip from wild-type (black), heterozygous (darker green), and homozygous *Myo15*$^{\Delta N/\Delta N}$ (bright green) littermates cultured in control conditions for 4 hours, from the data shown in Figure 3D. While the diameters of second row stereocilia were not statistically different between genotypes in OHC (A), the second row stereocilia from heterozygous and homozygous IHC were significantly thinner than in the wild-type littermates (C). Data are shown as mean ± SE. Statistical differences were analyzed using a linear mixed model. Statistical significance is shown as: *P<0.01, **P<0.001, ****P<0.0001; n.s., not significant. **(B,D)** Superimposed contours of OHC **(B)** and IHC **(D)** second row stereocilia tips from wild-type (*left*), heterozygous (*middle*), and homozygous *Myo15*$^{\Delta N/\Delta N}$ (*right*) littermates cultured in control conditions (*top*) or with 300 µM tubocurarine (*bottom*) for 4 hours. All contours were aligned to the tips of stereocilia, and the tip positions match the stereocilia height mean values shown in Figure 3C. Notice that, after 4 hours of MET channel blockage, wild-type stereocilia exhibit stereocilia tip thinning but have not shortened yet, heterozygous stereocilia exhibit more prominent thinning and some shortening, and homozygous stereocilia have even more exaggerated remodeling that has passed the initial thinning phase and results in larger shortening. For all panels, the age of explants is P7.

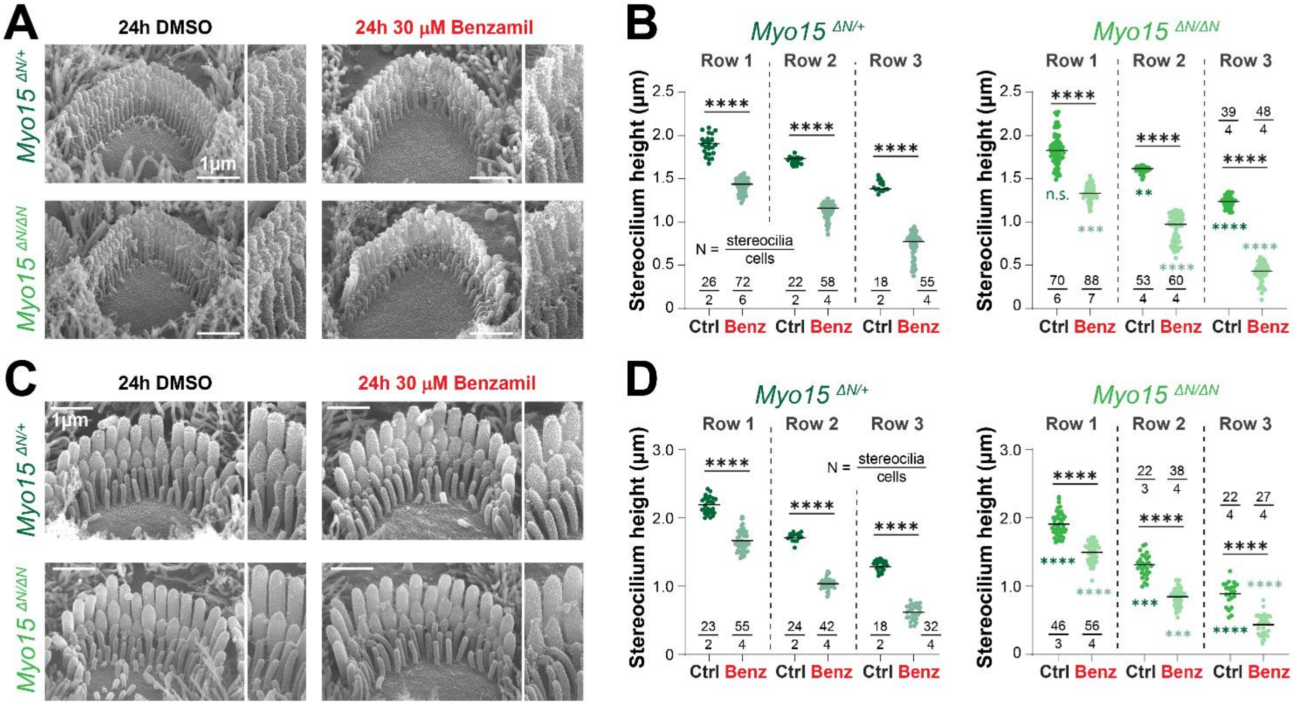


**Supplementary Figure 6. Exaggerated MET-dependent stereocilia cytoskeleton remodeling in the absence of the long isoform of MYO15A. (A,C)** Representative SEM images of OHC **(A)** and IHC **(C)** stereocilia bundles from heterozygous (*top*) and homozygous *Myo15*$^{\Delta N/\Delta N}$ (*bottom*) littermates cultured for 24 hours in vehicle control conditions (DMSO, *left*) or with 30 µM benzamil (*right*). **(B,D)** Heights of stereocilia from different rows of OHC **(B)** and IHC **(D)** bundles from heterozygous (darker shades of green) and homozygous *Myo15*$^{\Delta N/\Delta N}$ (brighter shades of green) cultured for 24 hours in vehicle control conditions (darker points) or in the presence of benzamil (lighter points). Data are from a single series of experiments. Horizontal lines indicate the mean. **S**tatistical significance is shown as: *P<0.05, **P<0.01, ***P<0.001, ****P<0.0001; n.s., not significant, from a Šídák's multiple comparisons test. Comparisons are made within the same genotype (in black) or between heterozygous and homozygous of the same stereocilia row and treatment (in green). Age of explants: P4 + 24 hours of incubation.

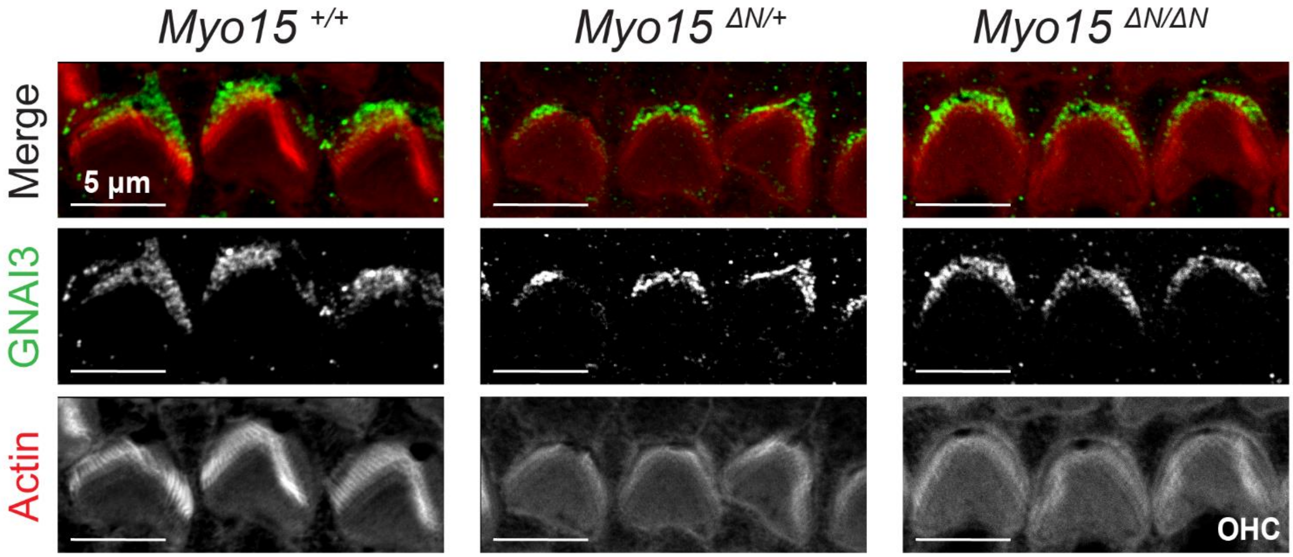


**Supplementary Figure 7. Localization of GNAI to the bare zone of OHC is maintained in *Myo15*$^{\Delta N/\Delta N}$ mice**. Maximum intensity projections of confocal stacks of OHC stereocilia (near the cuticular plate and ignoring stereocilia tips from the tallest row) from wild-type (*left*), heterozygous (*middle*), and homozygous *Myo15*$^{\Delta N/\Delta N}$ (*right*) mice immunolabeled against GNAI (green) and counterstained against F-actin with fluorescently-labeled phalloidin (red). Age of explants is P7.
